# DAF-2/insulin IGF-1 receptor regulates motility during ageing by integrating opposite signaling from muscle and neuronal tissues

**DOI:** 10.1101/2021.10.15.464514

**Authors:** Charline Roy, Laurent Molin, Allan Alcolei, Mathilde Solyga, Benjamin Bonneau, Camille Vachon, Jean-Louis Bessereau, Florence Solari

**Affiliations:** Université de Lyon, Université Claude Bernard Lyon 1, CNRS UMR 5284, INSERM U 1314, Institut NeuroMyoGène, Lyon 69008, France; Institut Curie, PSL Research University, Université Paris-Saclay, CNRS UMR 3347, INSERM U1021, Orsay 91405, France

**Keywords:** *daf-2*, motility, lifespan, DAF-16/FOXO, oxidative stress, mitochondria, insulin/IGF1 signaling, UNC-120/SRF

## Abstract

During ageing, preservation of locomotion is generally considered an indicator of sustained good health, in elderlies and in animal models. In *C. elegans*, mutants of the insulin-IGF-1 receptor DAF-2/IIRc represent a paradigm of healthy ageing, as their increased lifespan is accompanied by a delay in age-related loss of motility. However, these animals are less mobile than wild-type animals in early adulthood. Here we investigated the DAF-2/IIRc- dependent relationship between longevity and motility using an auxin-inducible degron to trigger tissue-specific degradation of endogenous DAF-2/IIRc. As previously reported, inactivation of DAF-2/IIRc in neurons or intestine was sufficient to extend the lifespan of worms, whereas depletion in epidermis, germline or muscle was not. However, neither intestinal nor neuronal depletion of DAF-2/IIRc prevented the age-related loss of motility. In 1-day-old adults, DAF-2/IIRc depletion in neurons reduced motility in a DAF-16/FOXO dependent manner, while muscle depletion had no effect. By contrast, DAF-2 depletion in the muscle of middle-age animals improved their motility independently of DAF-16/FOXO but required UNC-120/SRF.

Yet, neuronal or muscle DAF-2/IIRc depletion both preserved the mitochondria network in ageing muscle. Overall, these results show that the motility pattern of *daf-2* mutants is determined by the sequential and opposing impact of neurons and muscle tissues and can be dissociated from the regulation of the lifespan. This work also provides the characterization of a versatile tool to analyze the tissue-specific contribution of insulin-like signaling in integrated phenotypes at the whole organism level.

## INTRODUCTION

Ageing has long been regarded as an inevitable process resulting from a gradual and passive deterioration over time, as illustrated by the ‘wear and tear’ theory of ageing. However, genetic studies in the nematode *Caenorhabditis elegans* have revealed that a single mutation in the *daf-2* gene can double lifespan (Kenyon et al. 1993). *Daf-2* encodes the sole ortholog of the mammalian insulin and IGF-1 receptors (IIR) and its role in regulating lifespan appears to be conserved in mammals (Kenyon 2010). *Daf-2* mutants were initially isolated for their constitutive formation of diapausal larvae, called dauer (Daf-c phenotype). The long-lived and Daf-c phenotypes of *daf-2* mutants require the FOXO transcription factor DAF-16 (Ogg et al. 1997; Lin et al. 1997). Activation of the DAF-2/IIRc triggers a cascade of phosphorylation responsible for the retention of DAF-16/FOXO into the cytoplasm (Lin et al. 2001; Henderson and Johnson 2001; Lee et al. 2001; Li et al. 2021). Downregulation of this pathway promotes DAF-16/FOXO accumulation into the nucleus where it activates hundreds of genes (Tepper et al. 2013).

Decline in physical performance is a universal feature of ageing. *daf-2* mutants show improved motility in mid-adulthood that may be due to preserved motoneurons function (Liu et al. 2013; Hahm et al. 2015). Thus, *daf-2* mutants not only have a long lifespan, but are also considered healthier in old age. Yet, we and others (Huang et al. 2004; Mulcahy et al. 2012; Hahm et al. 2015; Podshivalova et al. 2017; Newell Stamper et al. 2018) noticed that in early adulthood, *daf-2* mutants are less mobile as compared to wild-type animals. This observation questions the relationship between lifespan and the other phenotypic traits considered as markers of good health.

To date, very little is known about the physiological expression pattern of DAF-2/IIRc. An immunolabelling approach has shown that DAF-2/IIRc is expressed mainly in the nervous system and in a pair of cells called “XXX cells” in the head and also in the epidermis (Kimura et al. 2011). In addition, two studies analyzed the requirement of DAF-2/IIR or DAF-16/FOXO in different tissues for lifespan regulation using transgene rescue of *daf-2 or daf-2; daf-16* mutants (Wolkow et al. 2000; Libina et al. 2003). The first study concluded that DAF-2 acts primarily in the nervous system, consistent with the expression profile of DAF-2/IIRc, while DAF-16/FOXO was later shown to act principally in the gut. This was counter-intuitive because these two proteins function in the same signaling pathway. The paradox was resolved by proposing that intestinal DAF-16/FOXO may trigger a secondary signal from the gut to induce inhibition of DAF-2/IIRc in distant neuronal tissues (Libina et al. 2003). However, these studies were based on standard transgenic strategies, the only tools available at the time, which lack some of the regulatory elements present in the endogenous loci and also lead to the overexpression of proteins encoded by the transgenes. New technologies have flourished to manipulate the endogenous expression of specific proteins, providing a way to re-examine the tissue-specific activities of DAF-2/IIRc, as we still do not know in which tissue wild-type DAF-2/IIRc functions for the regulation of the different phenotypes observed in *daf-2* mutants.

In this work, we investigated when and where DAF-2/IIRc is required to maintain worm motility during adulthood and how this phenotype relates to the dauer, lifespan and oxidative stress resistance phenotypes. In order to deplete DAF-2/IIRc protein in a spatially and temporally controlled manner, we generated alleles for conditional degradation by inserting an auxin-inducible degron (AID) (Zhang et al. 2015) and a fluorescent tag into the *daf-2* locus and constructed several independent strains to induce DAF-2/IIRc degradation in all cells or in neurons, muscle, intestine, germline or hypodermis. A previous study used similar tool without fluorescent tag to address the role of DAF-2/IIRc activity in neurons and intestine in the control of lifespan, dauer and oxidative stress resistance. Our work further explored the involvement of muscle, germline, hypodermis and combinations of tissues in these phenotypes and more specifically investigated the regulation of motility, whose maintenance is commonly regarded as a characteristic of healthy ageing.

Our results showed that DAF-2/IIRc is ubiquitously expressed in worms and can be efficiently degraded by the AID system. Degradation of DAF-2/IIRc in all tissues, from adulthood onwards, reproduced the lifespan and motility phenotypes of the reference *daf-2(e1370)* allele and the constitutive dauer phenotype in the progeny. Depletion of DAF-2/IIRc in neurons or in the gut was sufficient to extend the lifespan of the worms and only intestinal inactivation of DAF-2/IIRc reproducibly increased resistance to oxidative stress. Still, neither neuronal nor intestinal inactivation improved worm motility in adulthood. Neuronal depletion of DAF-2/IIRc unexpectedly downregulated motility from early adulthood, in a DAF-16/FOXO dependent manner. In contrast, muscle depletion was sufficient to improve motility in middle-aged worms, without affecting lifespan or resistance to oxidative stress. Finally, muscle inactivation of DAF-2/IIRc induced nuclear accumulation of DAF-16/FOXO in cells but did not require its activity for the regulation of motility which relies on the transcription factor UNC-120/SRF.

## RESULTS

### 1 The AID:mNeonGreen-tagged DAF-2 protein is functional and efficiently downregulated in the presence of auxin

The mNeonGreen (mNG) and degron sequences were added to the 3’ end (before the STOP codon) of the endogenous *daf-2* locus *(daf-2::AID::mNG or kr462)* so that DAF-2 degradation could be monitored via the loss of mNG fluorescence. A previous report used a similar strategy but without the fluorescent tag (Venz et al. 2021). DAF-2::AID::mNG was detected in head neurons, XXX cells and epidermis as reported earlier (Kimura et al. 2011) but also in the majority of the worm tissues, from the two-cell stage embryo (**Figure 1a**). *Daf-2(kr462)* worms were crossed with two independent lines that expressed in all tissues the plant ubiquitin ligase substrate recognition subunit, TIR1, an essential component of the AID system (see **Table S1** for strain description). We then verified that neither the tag nor the presence of the TIR1 transgene interfered with DAF-2 function. In the absence of auxin, *daf-2(kr462)* and *daf-2(kr462); Pubiquitous::TIR1* worms exhibited the same lifespan (**Figure 1b,c**; **Table S2**) and motility (**Figure 1d,e)** as wild-type worms. Furthermore, they did not enter the dauer stage at any temperature in the presence of food (**Table S3**), in contrast to the heat-sensitive reference *daf-2(e1370)* mutant*s* that showed a fully penetrant dauer constitutive phenotype. Thus, the addition of the degron and mNG sequences to the *daf-2* locus or TIR1 ubiquitous expression does not seem to impair DAF-2 function.

**Figure 1:**
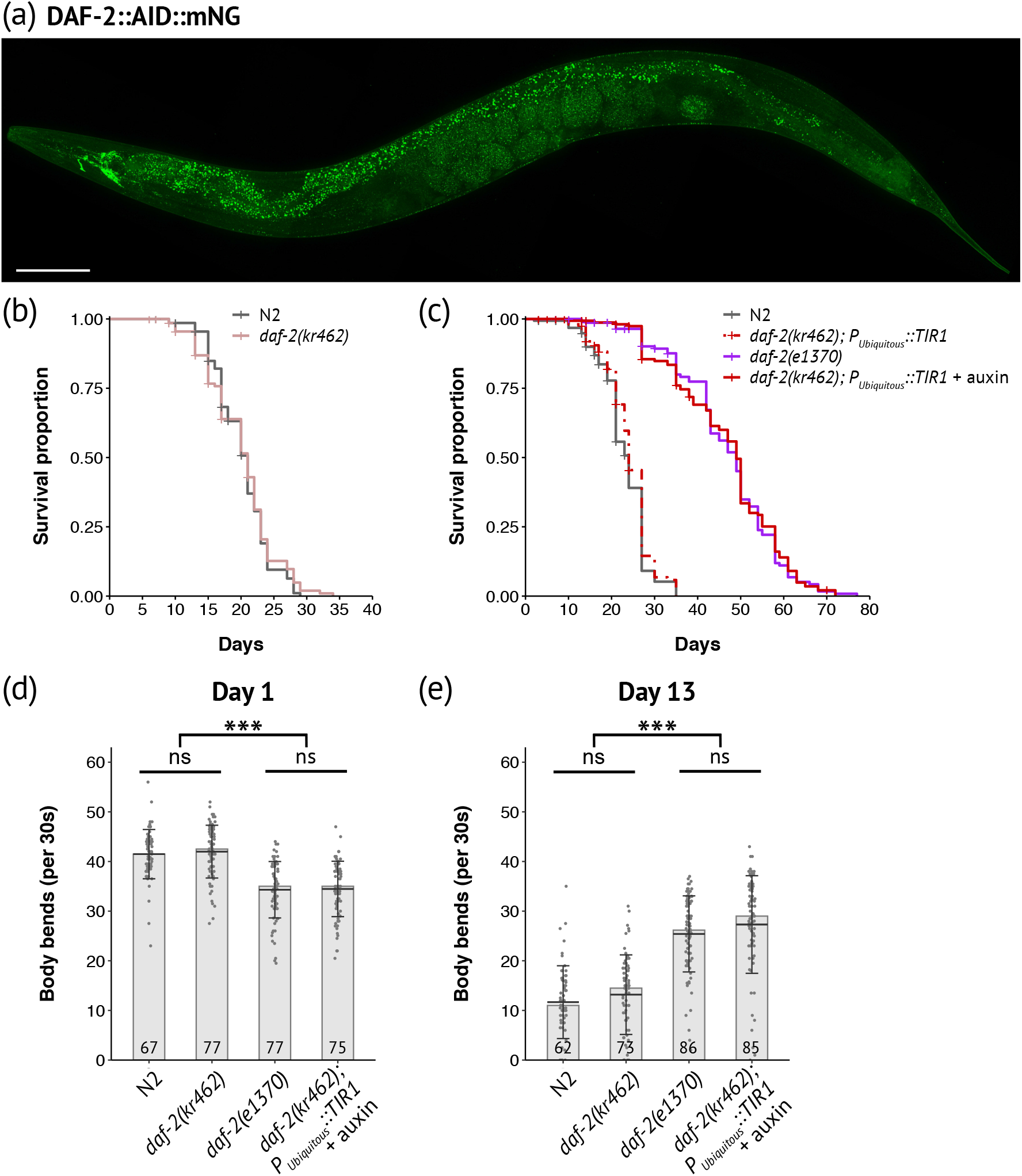
Expression pattern of DAF-2::AID::mNG and functional validation of its degradation by the auxin-inducible system. **(a)** Image of DAF-2::AID::mNG in 1-day-old *daf-2(kr462)* adult. Scale bar: 100 μm. **(b)** Survival curves of control (N2) and *daf-2(kr462)* animals (N= 2, n= 141 and 138 for N2 and *daf-2(kr462)*, respectively). **(c)** Survival curves of N2, *daf-2(e1370)* mutants, and worms with ubiquitous depletion of DAF-2. Data have been pooled from two independent experiments (n=155-160 for each genotype) in which two different *Peft-3::*TIR1 containing strains were tested. See **Table S1** for strain description and Table S2 for detailed lifespan data, replicates and statistics. **(d,e)** Body bends frequency at day 1 **(d)** and day 13 **(e)** of adulthood of N2, *daf-2(e1370)* or *daf-2(kr462)* worms, with or without ubiquitous expression of TIR1. The number of animals scored is indicated in each bar and corresponds to the pool of two experiments (see **Figure 4** for more replicates). Bars indicate median values, means are represented by black horizontal lines and brackets show standard deviations, ns: non-significant, ***: p < 0.001, Kruskal-Wallis and Dunn’s post hoc test with FDR method for adjusting p-value. All experiments were performed at 20°C.

In the presence of auxin, the fluorescence signal was strongly downregulated, confirming the efficiency of auxin-induced DAF-2 degradation (**Figure 2 and S1a**). We then assessed the dauer, lifespan and mobility phenotypes of *daf-2(kr462)* worms expressing TIR1 in all tissues compared to *daf-2(e1370)* mutants. All transgenic worms placed on auxin from hatching entered the dauer stage at 15, 20 or 25°C, (**Table S3** and Venz et al. 2021) mimicking the fully penetrant dauer phenotype of *daf-2(e1370)* mutants raised at the restrictive temperature of 25°C. Furthermore, when *daf-2(kr462)* worms were placed on auxin plates at a later developmental stage (L4) to bypass the dauer arrest, their lifespan was doubled, as *daf-2(e1370)* worms at 20°C (**Figure 1c and Table S2**). Remarkably, the ubiquitous degradation of DAF-2 also recapitulates the age-dependent motility phenotype of *daf-2(e1370)* mutants. Indeed, 1-day-old and 13-day-old *daf-2(kr462)* worms expressing TIR1 in all tissues showed a lower and higher frequency of body bends, respectively, compared to control animals of the same age (**Figure 1d,e**). Overall, these data demonstrate that the downregulation of DAF-2::AID::mNG correlates with a significant reduction of DAF-2 function, validating our experimental approach. However, the reference *daf-2(e1370)* allele is considered hypomorphic because predicted null *daf-2* mutations cause early developmental phenotypes with embryonic arrest (Gems et al. 1998). This suggests that some DAF-2 protein persists, consistent with the low level of fluorescence still detected in the presence of auxin (**Figure 2**). Nevertheless, our results show that inactivation of DAF-2 during development is not a prerequisite for lifespan extension, in agreement with previous results (Dillin et al. 2002; Venz et al. 2021) and that down-regulation of DAF-2 in adulthood is also sufficient to recapitulate the effect of constitutive down-regulation of DAF-2 on worm motility during ageing.

**Figure 2:**
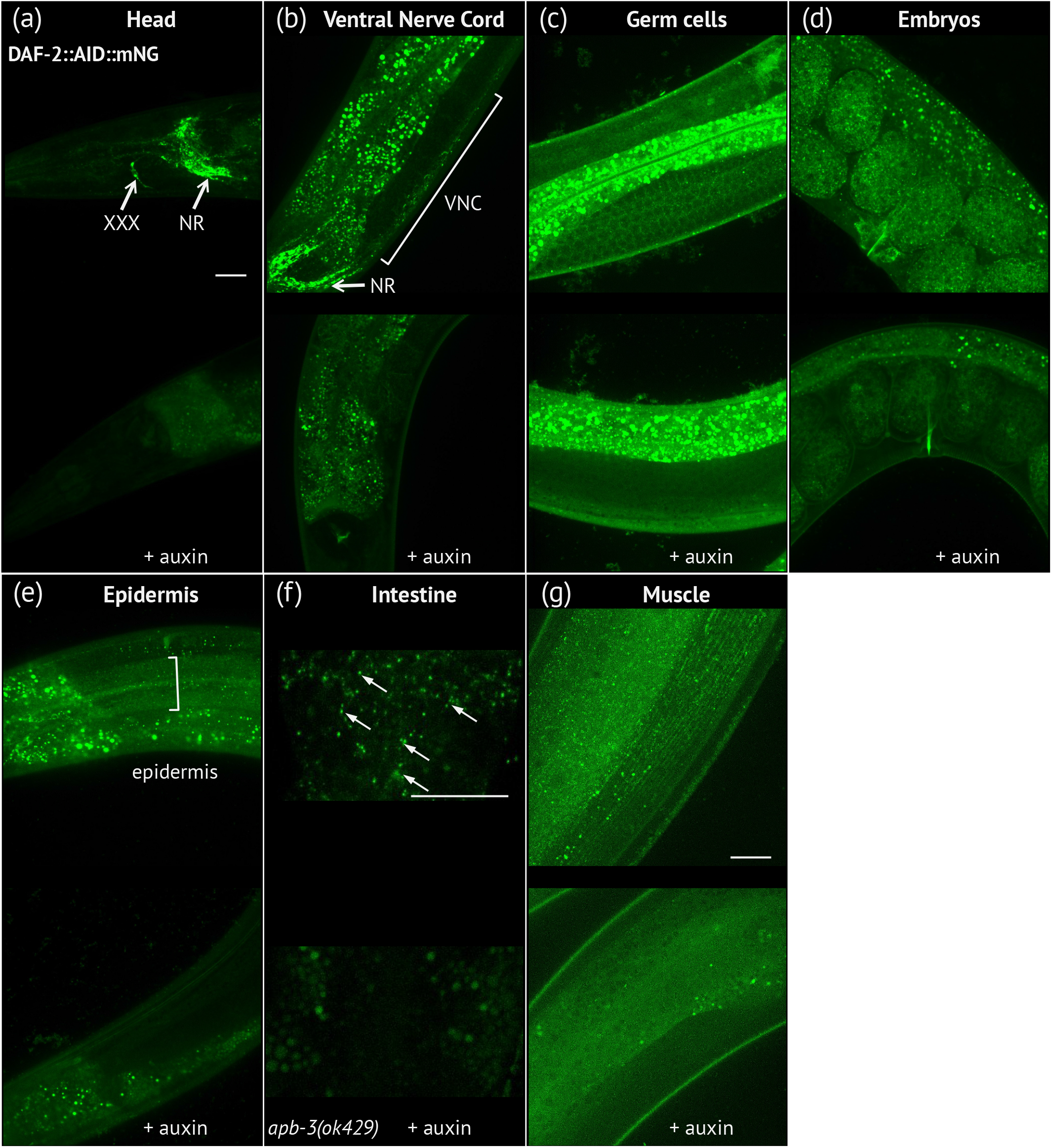
DAF-2::AID::mNG is effectively downregulated in the presence of ubiquitously expressed TIR1 after auxin treatment. **(a-g)** Images of DAF-2::AID::mNG in 1-day-old *daf-2(kr462)* adults expressing ubiquitous TIR1 and grown in the absence of auxin (upper panels) or after 24 hours of auxin treatment (lower panels). Images focus on specific body regions: the head **(a)**, showing strong expression in the nerve ring (NR) and the XXX cells; the neuronal cell bodies of the ventral nerve cord (VNC) **(b)**; the proliferating germ cells **(c)**; the embryos **(d)**; the epidermal syncytium **(e)**; the intestine **(f)** and the body wall muscles **(g)**. For the intestine **(f)**, images were taken in *apb-3(ok429)* mutant background in order to reduce unspecific intestinal autofluorescence (arrows indicate the specific DAF-2::AID::mNG associated signal). In all images, the remaining staining of the gut after auxin treatment corresponds to nonspecific autofluorescence that varies between animals. Similar results were obtained in 7-day-old animals (data not shown). Scale bars: 20 μm.

### 2 Intestinal and neuronal DAF-2 activities cooperate to regulate lifespan but are differently required for the resistance of worms to oxidative stress

We first examined the tissue-specific contribution of DAF-2 inactivation to the regulation of dauer and lifespan. Transgenic lines were generated to express TIR1 in muscle *(Pmyo-3)*, hypodermis *(Pdpy-17)*, neurons *(Prab-3)*, gut *(Pges-1)* or germline *(Psun-1)*. In addition to the previously described TIR1 transgenes, we generated new transgenes in order to test two independent lines for each tissue and thus limit potential confounding effect of the genetic background (see Experimental procedures and **Table S1**). In the presence of auxin, these TIR1 transgenes allowed efficient degradation of the DAF-2::AID::mNG protein in individual tissues, as indicated by the loss of fluorescent signals in young and middle-aged adults after 24 hours (**Figure S1b-h**). Dauer formation could not be achieved after degradation of DAF-2 in one given tissue (**Table S3**) thus suggesting that DAF-2 functions in several tissues and/or in cells other than the main tissues tested to control dauer entry, as also shown by Venz et al. (2021). These observations are in contradiction with previous data showing that overexpression of *daf-16* in the nervous system of *daf-2; daf-16* double mutants recapitulated the Daf-c phenotype of *daf-2* mutants (Libina et al. 2003). However, recent work on the tissue-specific activities of DAF-16 in a wild-type context argues that DAF-16 is required in several tissues to control the dauer phenotype (Aghayeva et al. 2021), in agreement with our results with DAF-2.

Inactivation of DAF-2 in muscle, hypodermis or germline did not reproducibly affect lifespan suggesting that depletion of DAF-2 in these tissues may not be sufficient to impact this phenotype (**Figure 3a-f; Figure S2d-g and Table S2**). However, we cannot completely rule out that a small amount of undegraded DAF-2 remained in these tissues.

**Figure 3:**
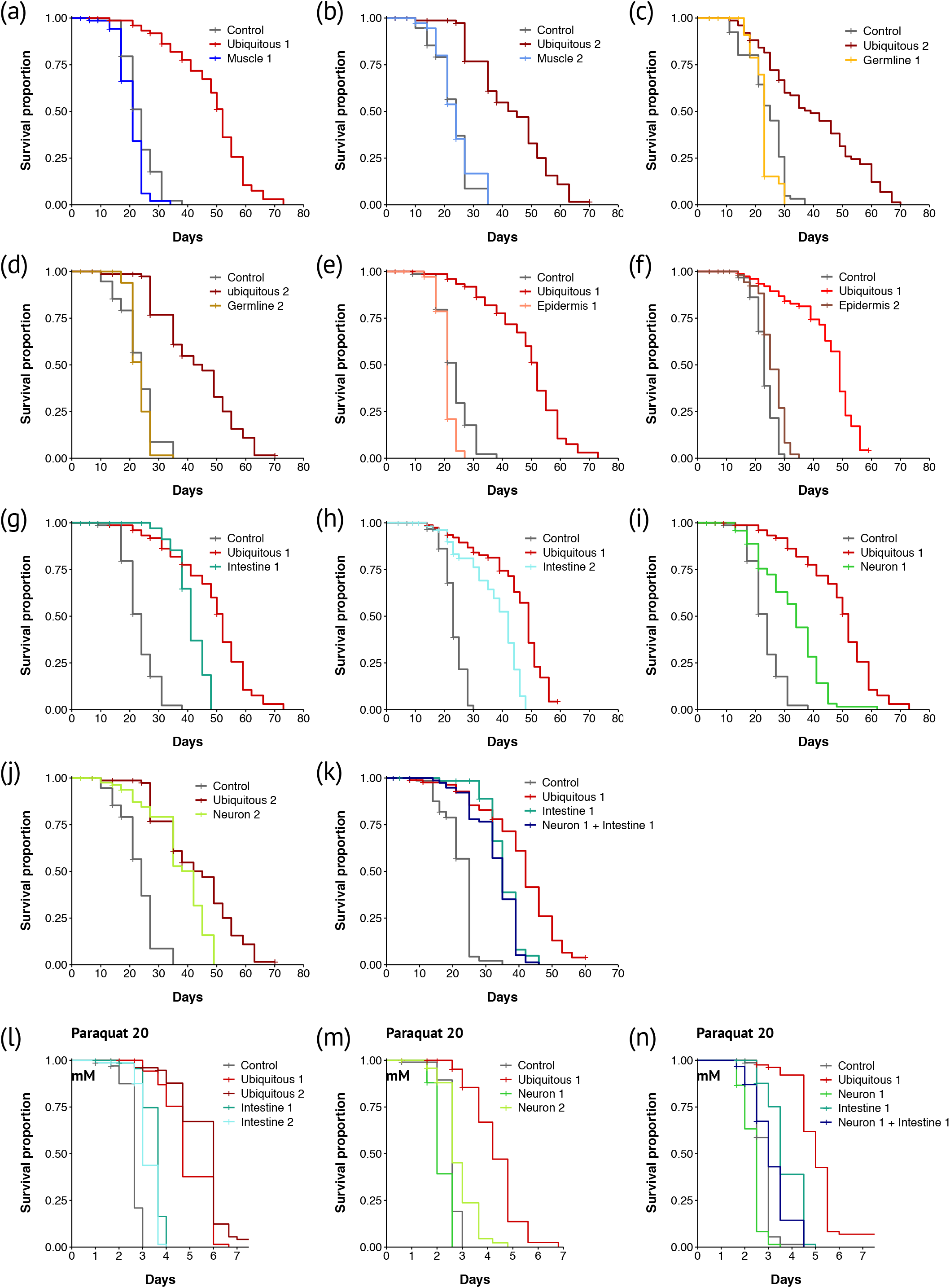
Inactivation of DAF-2 in neurons or in the gut is sufficient to increase lifespan. **(a-k)** Survival curves of animals with DAF-2 depletion in all cells **(a-k)**, muscle **(a-b)**, germline **(c-d)**, epidermis **(e-f)**, intestine **(g-h)**, neurons **(i-j)** or neurons and intestine **(k)**. Numbers (ex: Muscle 1, Muscle 2) refer to distinct alleles driving TIR1 expression (See Table S1 for complete strain description). For each condition, n is about 80 individuals. Some experiments were split in separate graphs for clarity, thus some graphs share the same negative and positive controls: **(a)**, **(e)**, (g), **(i)**; **(b)**, **(d), (j)**; **(f)** and **(h)**. The control conditions correspond to the N2 or *daf-2(kr462)* strains, in the presence of EtOH or auxin, whose lifespans did not show significant differences. **(l-n)** Survival curves of animals with DAF-2 depletion in all cells (**l-n**), intestine (**l**), neurons (**m**) or neurons and intestine (**n**) in presence of 20 mM paraquat. Controls correspond to *daf-2(kr462)*. All strains including controls have been treated with auxin. For each condition, 75 to 100 individuals have been assayed. For detailed lifespan data, replicates, statistics, and summary of independent assays see Table S2 and Figure S2.

Degradation of DAF-2 in neurons or intestine was sufficient to increase the median lifespan by 41% and 57,1% respectively. However, none of these tissues recapitulated the lifespan increase caused by ubiquitous DAF-2 inactivation (**Figure 3g-j; Figure S2h, i**). Combined DAF-2 degradation in both neurons and gut did not further extend the lifespan of animals as compared to worms with intestinal or neuronal DAF-2 degradation alone **(Figure 3k; Figure S2j).** Thus, additional cells or a different combination of tissues may be involved in the regulation of lifespan by DAF-2.

Resistance to oxidative stress has been proposed as a mechanism responsible for the extension of lifespan by DAF-2 (Honda and Honda 2002). In the presence of paraquat, a reactive oxygen species generator, ubiquitous or intestinal inactivation of DAF-2 consistently increased the lifespan of worms compared to control worms, although to a lesser extent for intestinal lines (**Figure 3l; Figure S2k,l,n**). Neuronal inactivation of DAF-2 reproducibly resulted in a slight increase in paraquat resistance in one line, and a slightly lower resistance in the second (**Figure 3m, Figure S2l, m, n**), although both lines were long-lived in the absence of paraquat (**Figure 3i, j).** In addition, neuronal inactivation of DAF-2 reduced the resistance to oxidative stress of worms with intestinal DAF-2 depletion **(Figure 3n, Figure S2n**), but did not shorten their lifespan **(Figure 3k**). These data suggest that neuronal and intestinal inactivation of DAF-2 triggers lifespan extension mechanisms that are at least partially independent of oxidative stress resistance.

### 3 DAF-2 neuronal signaling is required for worm motility in early adulthood, while DAF-2 muscle signaling impairs motility from mid-adulthood

In order to further characterize the fitness of long-lived worms we measured their body-bend frequency in liquid medium (BBF), as a proxy of physical performance (Duhon and Johnson 1995; Laranjeiro et al. 2019). The motility of worms with DAF-2 inactivation in the intestine was similar to that of control worms on days 1 and 13 of adulthood (**Figure 4a-c).** Thus, the signaling from the intestine upon DAF-2 degradation is sufficient to prolong lifespan, but does not markedly affect the function of the neuromuscular system with age.

**Figure 4:**
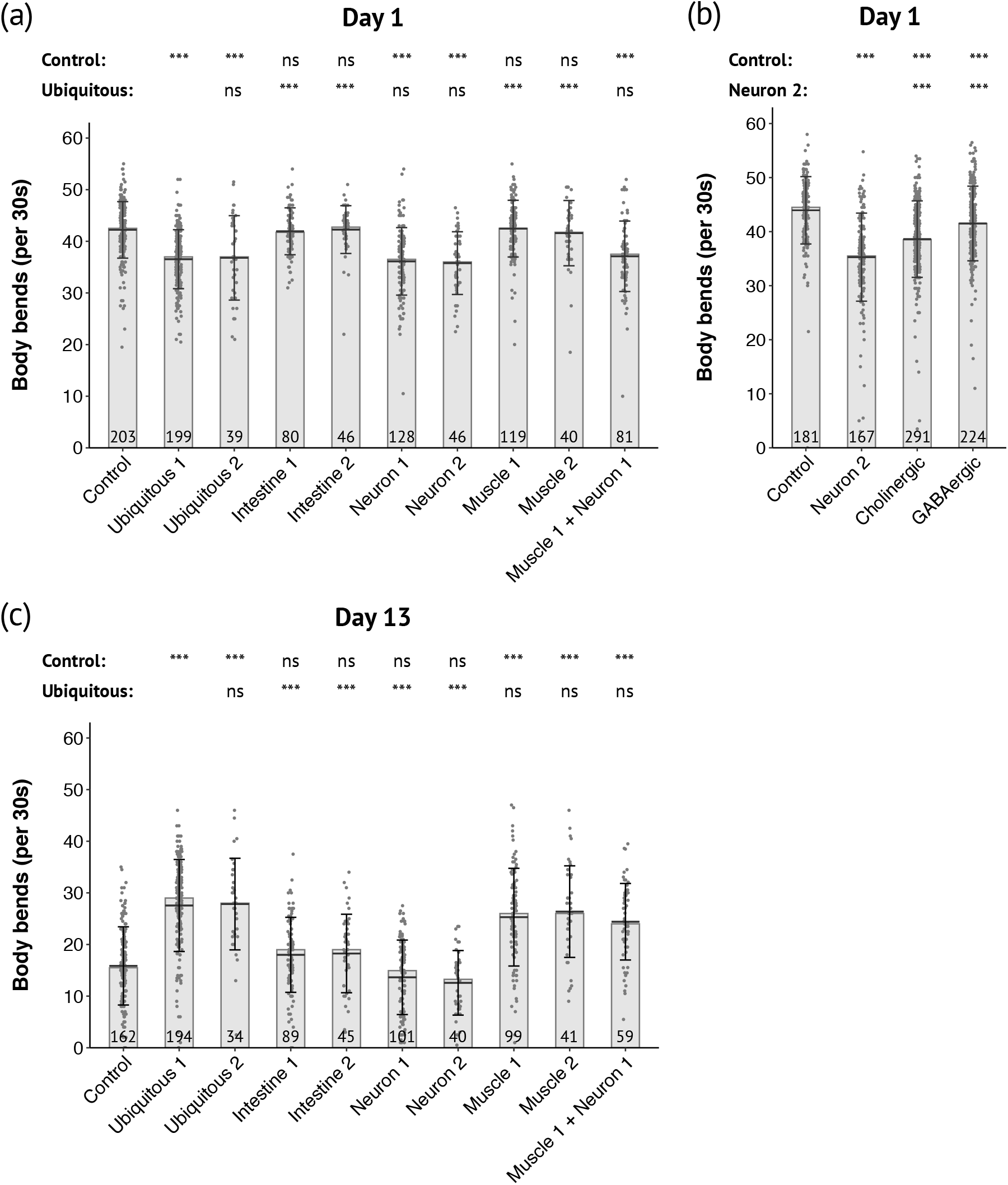
DAF-2 degradation in neurons or muscles differentially alters mobility in an age-dependent manner. **(a-c)** Body bends frequency of 1-day-old **(a-b)** and 13-day-old **(c)** adults with depletion of DAF-2 in all cells or in the intestine, muscle, neurons or muscle and neurons as indicated. Control corresponds to *daf-2(kr462)* in presence of auxin. Numbers (ex: Muscle 1, Muscle 2) refer to distinct alleles driving TIR1 expression (See Table S1 for the description of strains). **(a and c)** Pooled data from 5 independent experiments. Each replicate included the control strain and several tissue specific strains. **(b)** Pooled data from 7 independent experiments in which were assayed 4 and 3 independent lines for depletion of DAF-2 in cholinergic or GABAergic neurons, respectively. The number of animals scored is indicated in each bar. The bars correspond to the median values, the means are represented by black horizontal lines and brackets show standard deviations. Comparisons were done with Kruskal-Wallis, Dunn post-hoc tests with FDR method to adjust p-value, ns: not significant, ***: p_adjusted_ < 0.001. Statistics are presented as two lines that include comparison with the control strain or with one specific strain (ubiquitous 1 for **(a and c)** and neuron 2 for **(b)**).

Neuronal inactivation of DAF-2 significantly reduced the worm’s BBF of 1-day-old animals and mirrored the motility phenotype of animals with whole-body degradation of DAF-2 **(Figure 4a).** This result was unexpected, as Liu et al. had previously reported that the *daf-2* mutation delays the functional decline of neurons at the neuromuscular junction (NMJ), which begins on day five of adulthood in wild-type worms (Liu et al. 2013). We thus assessed cholinergic neurotransmission in our strains by treating worms with aldicarb. Aldicarb is an inhibitor of acetylcholinesterase, which induces worm paralysis due to the accumulation of acetylcholine in the synaptic cleft. Consistent with Liu et al. data, we observed an increase in cholinergic neurotransmission, indicated by an accelerated paralysis in response to aldicarb, when DAF-2 was ubiquitously inactivated in middle-aged animals but also in 1-day-old animals (**Figure S3**). However, this phenotype was uncoupled from the motility phenotype since worms with neuronal depletion of DAF-2 behaved on aldicarb like control worms **(Figure S3).**

The regulation of worm motility relies on a complex neuronal network that involves different class of interneurons, excitatory cholinergic and inhibitory GABAergic motoneurons (Zhen and Samuel 2015). Previous report showed that DAF-2 is expressed in both cholinergic and GABAergic neurons (Taylor et al. 2021). Depletion of DAF-2 in either cholinergic or GABAergic neurons only was sufficient to impede worm’s mobility in 1-day old animals thus suggesting that DAF-2 functions in both types of neurons to control motility in young adults **(Figure 4b)**.

In contrast to neuronal DAF-2, muscle inactivation of DAF-2 did not affect the BBF of 1-day-old adult animals but was sufficient to recapitulate the higher BBF of 13-day-old adult animals with ubiquitous DAF-2 depletion **(Figure 4a, c)**, although it affected neither lifespan **(Figure 3a, b; Figure S2d)** nor resistance to oxidative stress **(Figure S2o, p).** Furthermore, inactivation of DAF-2 in neurons did not suppress the beneficial impact of muscle DAF-2 inactivation on motility (**Figure 4c**).

Taken together, these data support a critical role for wild type DAF-2 activity in muscle in negatively regulating motility in middle age, while neuronal DAF-2 promotes motility in early adulthood.

### 4 DAF-16 nuclear accumulation upon tissue-specific inactivation of DAF-2

Early studies on the tissue-specific activities of DAF-2 and DAF-16 have suggested that down-regulation of the DAF-2 signaling pathway in one tissue induces its inhibition in distant tissues (Wolkow et al. 2000; Libina et al. 2003). However, these results were obtained in a sensitized genetic background, as all cells were mutant for *daf-2*. More recent data supported this model in a *daf-2(+)* background by showing that overexpression of intestinal or neuronal DAF-16 induces nuclear accumulation of DAF-16::GFP in distant tissues (Uno et al. 2021). To avoid potential issues associated with reporter transgenes such as overexpression, we endogenously tagged DAF-16/FOXO with wrmSCARLET using CRISPR/Cas-9 mediated genome engineering. DAF-16:: wrmSCARLET was detected in all somatic tissues and the germline during adulthood, with highest expression in neurons, and localized mainly in the cytoplasm of all cells **(Figure 5a)**. Ubiquitous depletion of DAF-2 from the L4 stage induced nuclear accumulation of DAF-16 in all tissues **(Figure 5b)** and the whole progeny entered the dauer stage (data not shown), confirming that the DAF-16:: wrmSCARLET protein was functional.

**Figure 5:**
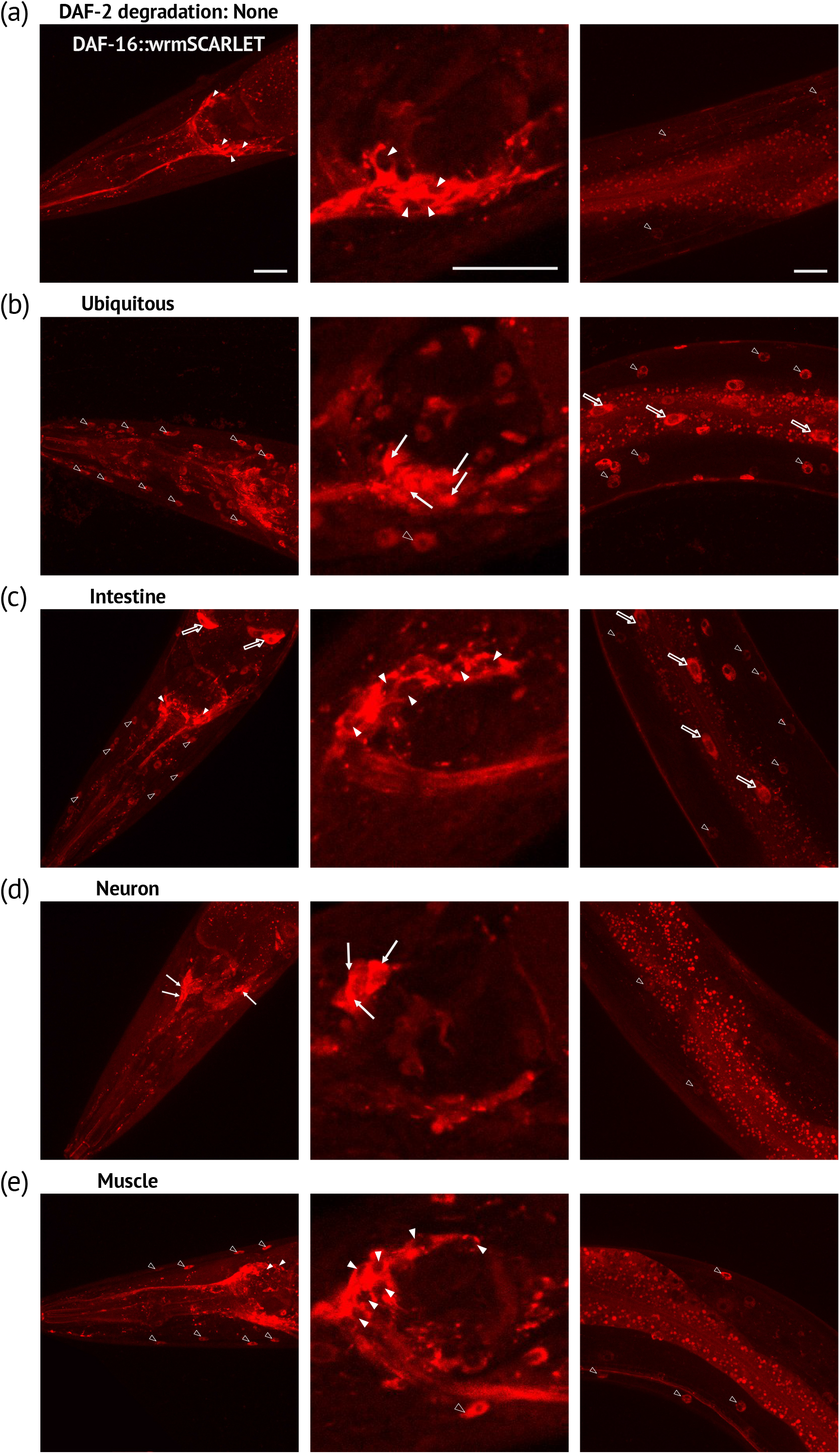
Regulation of DAF-16 subcellular localization by tissue-specific depletion of DAF-2. **(a-e)** Images of DAF-16::wSCARLET in the head (left and middle panels) or the anterior body (right panels) of 1-day-old *daf-16(kr535); daf-2(kr462)* adults without (a) or with ubiquitous (b), intestinal (c), neuronal (d) or muscular (e) depletion of DAF-2 caused by 24 hours of auxin treatment (b-e). Empty arrowheads, full arrows and empty arrows indicate muscle, neuron and intestinal nuclei respectively; full arrowheads correspond to the cytosol of neurons. For numbers and percentage of worms with a strong DAF-16::wSCARLET nuclear signal, see Table S4. Scale bars: 20 μm.

Depletion of DAF-2 in the intestine was associated with nuclear accumulation of DAF-16::wrmSCARLET in the gut and distant tissues that do not seem to include neurons (**Figure 5c**). In contrast, tissuespecific degradation of DAF-2 in neurons or muscle from L4 onwards induced nuclear accumulation of DAF-16:: wrmSCARLET in the same tissue but not in distant tissues **(Figure 5 d,e)**. This suggests that the regulation of motility by neuronal and muscle DAF-2 activity does not rely on DAF-2 inactivation in distant tissues.

### 5 The control of motility by the neuronal and muscle activities of DAF-2 requires DAF-16 and UNC-120 respectively

To investigate the functional significance of cell-autonomous regulation of DAF-16 nuclear accumulation by DAF-2, we crossed worms in which DAF-2 is inactivated in muscle or neuronal cells with worms carrying the *daf-16(ot853)* allele that encodes DAF-16::mNeonGreen::AID (Bhattacharya et al. 2019). *daf-16(ot853); daf-2(kr462)* worms showed a slight reduction in motility compared to control *daf-2(kr462)* worms (**Figure 6a)** Nevertheless, degradation of DAF-16 in worms with neuronal inactivation of DAF-2 did not decrease the motility of 1-day-old animals as observed when only DAF-2 is inactivated (**Figure 6a).** This data strongly suggests that inactivation of DAF-16 suppressed the motility phenotype associated to DAF-2 neuronal depletion. In contrast, the increase in motility of 13-day-old animals with muscle inactivation of DAF-2 was not affected by depletion of DAF-16 in the same tissue (**Figure 6b).** We then tested whether the transcription factor UNC-120/SRF was required, as we have previously shown that inhibition of *unc-120* expression in *daf-2(e1370)* mutants significantly reduced their motility from mid-adulthood (Mergoud dit Lamarche et al. 2018). Furthermore unc-120 is expressed in muscles, but not in neurons (Mergoud dit Lamarche et al. 2018). RNAi mediated inactivation of *unc-120* from adulthood in worms with muscle-specific inactivation of DAF-2 also suppressed DAF-2 impact on worm motility (**Figure 6c**).

**Figure 6:**
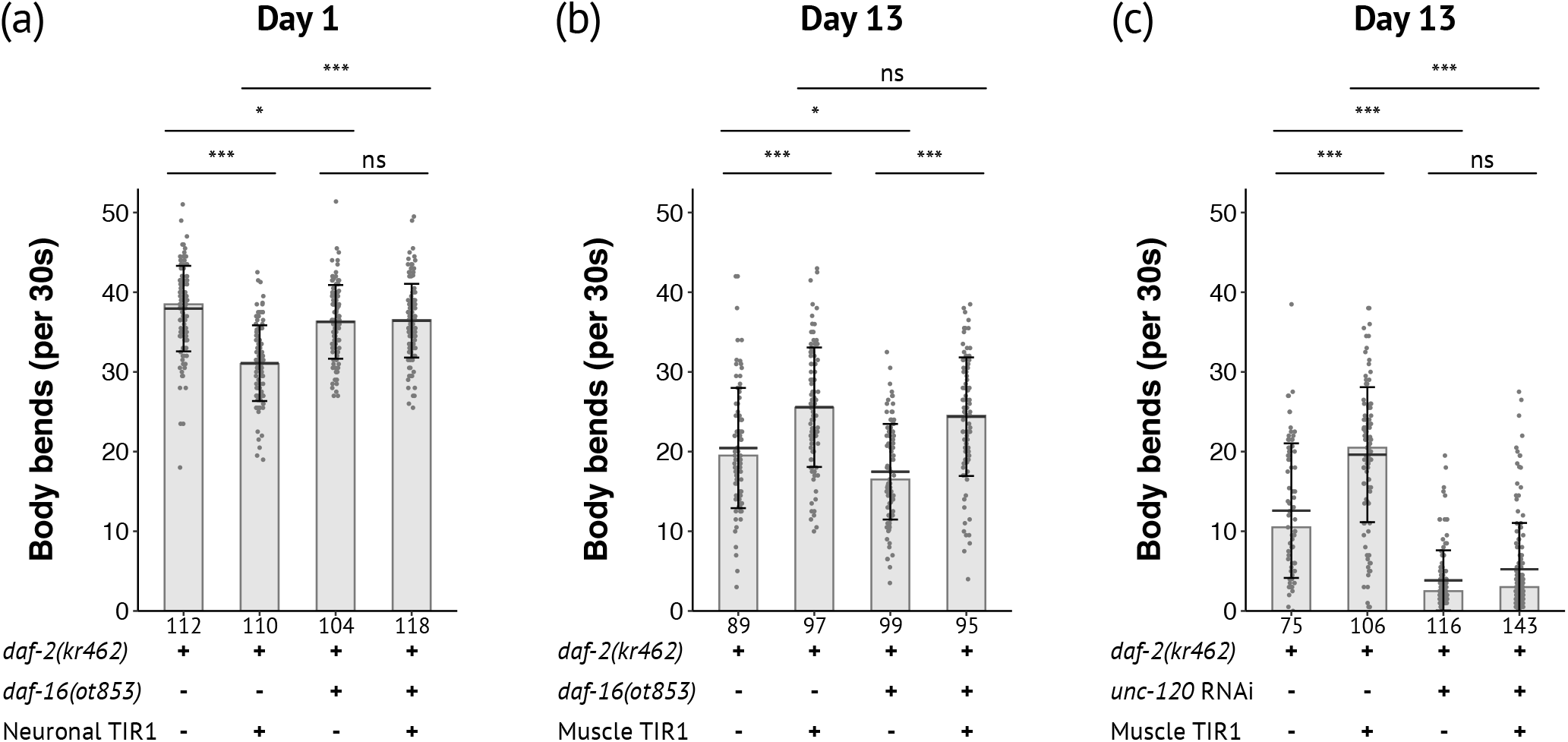
DAF-16 is required for motility regulation when DAF-2 is inactivated in neurons but not in muscles. Body bends frequency of 1-day-old **(a)** and 13-day-old **(b, c)** adults with depletion of DAF-2 and DAF-16 in neurons or muscle **(a, b)** or with downregulation of *unc-120* by RNAi in a *rrf-3(pk1426)* genetic background **(c)** (see experimental procedure and Table S1 for detailed genotype of strains). Data from three independent experiments were pooled. The number of animals scored is indicated under each bar. The bars correspond to the median values, the means are represented by black horizontal lines and brackets show standard deviations. Comparisons were done with Kruskal-Wallis, Dunn post-hoc tests with FDR method to adjust p-value, ns: not significant, *: p_adjusted_ < 0.05, ***: p_adjusted_ < 0.001

Overall, while DAF-16 accumulated in the nuclei of neurons or muscles after inactivation of DAF-2 in the same tissue, it is essential for the regulation of motility in neurons but not in muscles in which DAF-2 required UNC-120 for motility regulation.

### 6 Both neuronal and muscular DAF-2 depletion prevents muscle mitochondria fragmentation with age

We and others have previously shown that the morphology of muscle mitochondria changes from an interconnected to a fragmented network during muscle ageing and that this phenotype is associated with loss of motility in old age (Mergoud dit Lamarche et al. 2018). To further investigate the cell autonomous function of DAF-2 in muscle ageing, we monitored the muscle mitochondria pattern with age. We visualized mitochondria by expressing a single-copy insertion of a reporter encoding the N-terminal segment of TOM20 (responsible for its anchoring to the mitochondrial outer membrane) fused to the fluorescent protein wrmSCARLET, under the control of the muscle *pmyo-3* promoter (see Experimental procedures). It is noticeable that previously characterized worm strains carrying similar reporters but as an integrated multicopy array showed earlier mitochondrial fragmentation, probably due to overexpression of the fusion protein (Regmi et al. 2014a; Mergoud dit Lamarche et al. 2018). The ubiquitous degradation of DAF-2 delayed the fragmentation of mitochondria **(Figure S4)** in agreement with previous observations with the *daf-2(e1370)* mutant (Mergoud dit Lamarche et al. 2018). Muscle inactivation of DAF-2 was sufficient to prevent muscle mitochondria fragmentation. However, neuronal depletion of DAF-2 gave similar results, alone or in combination with muscle depletion of DAF-2 **(Figure S4).** Thus, inactivation of DAF-2 in muscles or neurons is sufficient to prevent muscle mitochondrial fragmentation, suggesting that DAF-2 acts both autonomously and non-autonomously to maintain muscle integrity during ageing.

## Discussion

In this work, we investigated the relationship between lifespan and motility phenotypes caused by DAF-2/IIRc inactivation by analyzing the tissue-specific functions of DAF-2/IIRc in the regulation of these phenotypes. To this end, we have created a reporter line that allows the visualization of both DAF-2/IIRc expression and its degradation by the TIR1 auxin-inducible system in live animals.

### 1 *Daf-2(kr462)* transgenic animals as a model to study DAF-2 expression and function

As in other species, multiple isoforms of DAF-2/IIRc have been described, including DAF-2A and DAF-2C, which differ by the size of the α-chain of the α2β2 tetramer, and are analogous to mammalian IRA and IR-B, respectively (Ohno et al. 2014). The mNeonGreen tag was inserted into the C-terminus shared by these two isoforms which control lifespan, dauer, heat tolerance (A and C) and avoidance behaviour (C only). Three shorter isoforms have been reported (Ohno et al. 2014). DAF-2B, which retains the extracellular ligand-binding domain but lacks the intracellular signaling domain, modulates insulin signaling by sequestering insulin peptides, but its expression is restricted to the developmental larval stages and is no longer observed in the adult stage (Martinez et al. 2020). The DAF-2D and E isoforms lack part of the β-chain and IRS1 interaction domain and their function remains to be defined. We showed that ubiquitous inactivation of DAF-2 A and C, from the pre-adult L4 stage, is sufficient to recapitulate the motility and longevity phenotypes of *daf-2(e1370)* mutants, which carry a single mutation in the DAF-2 kinase domain common to DAF-2A, C, D and E isoforms (Kimura et al. 1997). Thus, DAF-2A and/or DAF-2C play a major role in the regulation of motility, lifespan, dauer and oxidative stress resistance while the DAF-2D and E isoforms are not essential.

We detected DAF-2::AID::mNG receptor expression throughout the body, as in flies and mammals, and more extensively than previously appreciated using immunochemistry (Kimura et al. 2011). Interestingly, DAF-2::AID::mNG receptors fluorescence appeared as a punctate pattern in the cytoplasm of most cells, rather than enriched at the cell membrane. This pattern may highlight DAF-2/IIRc biosynthetic pathway and/or endosomal signaling compartment. Recent data obtained in mammalian cell culture also showed that the majority of insulin receptors are localized within intracellular vesicles under regular culture conditions (Boothe et al. 2016). Internalization of the insulin receptor is necessary to shut down insulin signaling, but it also induces endosome-specific signal transduction (Morcavallo et al. 2014). Increasing evidence suggests that alterations in the insulin receptor trafficking can lead to severe insulin resistance (Chen et al. 2019). *daf-2(kr462)* transgenic animals will be useful to further study the conservation of insulin/IGF-1 receptor biosynthesis and trafficking and its potential deregulation in the context of different *daf-2* mutants.

### 2 DAF-2/IIRc activity in neurons and intestine limits wild-type lifespan via shared mechanisms that do not involve inter-organ inactivation of the DAF-2/IIRc

Our data pointed to the importance of both intestine and neurons for lifespan regulation by the DAF-2/IIRc pathway as previously proposed (Wolkow et al. 2000; Libina et al. 2003; Uno et al. 2021; Venz et al. 2021). Moreover, we showed that combined DAF-2/IIRc degradation in the neurons and in the gut did not further extend the lifespan of the animals compared to worms with DAF-2/IIRc degradation in the gut. This suggests that DAF-2/IIRc signaling in the gut and neurons share downstream mechanisms for the regulation of lifespan. Surprisingly, although intestinal DAF-2/IIRc inactivation conferred resistance of animals to oxidative stress, the correlation between longevity and oxidative stress resistance phenotypes was not verified with neuronal inactivation of DAF-2/IIRc. The link between oxidative stress resistance and lifespan extension has been a matter of intense debate for years (Dues et al. 2017, 2019). Our data suggest that while neural and intestinal DAF-2/IIRc affect lifespan through common mechanisms, those mechanisms do not seem to involve resistance to oxidative stress. Also, we cannot exclude that our results might have been different if we had used another oxidative stressor, as we only studied a severe one (paraquat).

Shared mechanisms may result from the inactivation of DAF-2/IIRc in distant tissues (ie gut or neurons) when DAF-2/IIRc is inactivated in one tissue (ie neurons or gut respectively) as previously proposed (Libina et al. 2003; Uno et al. 2021). **(Figure 5).** This discrepancy is probably linked to the lines used for monitoring DAF-16 subcellular localization. Previous studies used wild-type animals carrying an integrated multicopy array *(Pdaf-16::daf-16::gfp)* that exhibit several phenotypes that are reminiscent of DAF-16 gain-of-function (Henderson & Johnson 2001). Therefore, gut and neuronal DAF-2/IIRc function in lifespan does not seem to depend on DAF-2/IIRc inactivation through inter-organ communication.

*Daf-2(e1370)* prevents loss of gut integrity (Gelino et al. 2016) and visceral pathologies associated with ageing that limits worm survival (Ezcurra et al. 2018). During the last years several longevity pathways have been shown to involve brain-gut communications in *C. elegans* (Prahlad et al. 2008; Durieux et al. 2011; Taylor and Dillin 2013; Berendzen et al. 2016; Shao et al. 2016; Zhang et al. 2018), which may be explored using tissue-specific inactivation of DAF-2/IIRc to better understand the role of DAF-2/IIRc in lifespan regulation.

### 3 Neuronal inactivation of DAF-2/IIRc downregulates motility in a DAF-16 dependent manner during early adulthood

Declined motility of worms with age has been associated with both decreased neural stimulation and the loss of muscle cell integrity (Mulcahy et al. 2012; Liu et al. 2013; Mergoud dit Lamarche et al. 2018). Based on electrophysiological data, Liu et al (2013) reported that synaptic transmission defects in motor neurons appear as early as day 5 of adulthood, while muscle contraction defects do not occur before day 11 of adulthood. They also showed that *daf-2(e1370)* mutation delayed the functional decline of neurons at the neuromuscular junction. Consistent with these data, we observed that whole-body inactivation of DAF-2 increased excitatory cholinergic neurotransmission at the neuromuscular junction in middle-aged animals, but also as early as day 1 of adulthood. (**Figure S3)**. However, when we further tested DAF-2 inactivation in muscle or neurons only, worms did not show this phenotype at neither day 1 or day 14 of adulthood (Figure S2). Those data strongly suggest that the motility phenotype associated to *daf-2* inactivation in either muscle or neurons does not rely on the modulation of cholinergic transmission at the neuromuscular junction, although we cannot rule out subtle defects that would only be apparent at physiological concentrations of acetylcholine.

Interestingly those results also show that the increase in neurosecretion in *daf-2* mutants (our results and Li et al., 2013) can be dissociated from their motility phenotype. The mechanisms responsible for this discrepancy remained to be elucidated.

### 4 Muscle mitochondria morphology is regulated both autonomously and non-autonomously by DAF-2 activity

Mitochondria are dynamic organelles that undergo cycles of fusion and fission, important for the maintenance of their membrane potential and for mitophagy, respectively (Kleele et al. 2021). Interconnected or elongated mitochondria exhibit increased efficiency of ATP production and reduced generation of reactive oxygen species (ROS). Conversely, fragmented morphology is associated with reduced ATP production and uncoupling of mitochondria. Aberrant morphology and mitochondrial dysfunction are universal features of ageing in invertebrates and vertebrates. In *C.elegans*, those changes may have a causal role in ageing as modulation of either fusion or fission affect both lifespan and motility of worms (Chaudhari and Kipreos 2017; Weir et al. 2017; Byrne et al. 2019; Liu et al. 2020). We and others have shown that muscle ageing is associated with progressive mitochondria fragmentation (Regmi et al. 2014b; Mergoud dit Lamarche et al. 2018) that is delayed in *daf-2(e1370)* mutants (Mergoud dit Lamarche et al. 2018).

Muscle inactivation of DAF-2/IIRc was sufficient to prevent fragmentation of mitochondria, suggesting that the increased motility of the worms relies on improved mitochondrial homeostasis. However, neuronal inactivation of DAF-2/IIRc protein resulted in a similar mitochondrial phenotype, but did not improve motility in middle age. Thus, although a young mitochondrial network may be a prerequisite for the maintenance of the neuromuscular system with ageing, this may not be sufficient for the upregulation of motility by muscle DAF-2/IIRc.

### 5 DAF-2 acts primarily in muscle to control motility in ageing worms

Muscle DAF-2/IIRc plays a major role in delaying the loss of worm motility observed in *daf-2* mutants, while it did not affect worm life span or resistance to oxidative stress. Furthermore, DAF-16/FOXO was dispensable for the regulation of motility by muscular DAF-2/IIRc while UNC-120/SRF was required. These observations agree with our previous data obtained in the context of *daf-2 (e1370)* mutants, and support the cell-autonomous impact of DAF-2/UNC-120 on worm motility in middle age.

Our results do not exclude the existence of a secondary signal from muscle to neurons that could improve motility. Indeed, previous work has identified the miRNA *mir-1*, as a regulator of a retrograde signal from muscle to neurons that modulates neuronal activity (Simon et al. 2008) and *mir-1* inactivation improves worm motility under pathological conditions (Schiffer et al. 2021). Interestingly, mammalian SRF has been shown to negatively regulate *mir-1* expression (Zhang et al. 2011; Tritsch et al. 2013). Identification of transcriptional targets of UNC-120/SRF in the context of muscle DAF-2 inactivation should help to better define the cellular mechanisms involved in the regulation of motility by muscle DAF-2.

Overall, we have developed and characterize a powerful tool to explore DAF-2 function with ageing, uncovering unexpected findings regarding tissue-specific roles of DAF-2 in the regulation of dauer, lifespan, resistance to oxidative stress, and motility, as well as in the cross-talk between tissues.

Although both DAF-2 and DAF-16 are ubiquitously expressed, our approach identifies a tissue-specific, antagonistic and age-dependent role for DAF-2/DAF-16 and DAF-2/UNC-120 signaling in the regulation of motility. Numerous phenotypes have been associated to *daf-2* mutants and the future challenge will be to define the contribution of the different tissue-specific activities of DAF-2 in the regulation of these phenotypes and their downstream effectors.

## Experimental Procedures

### *C. elegans* strains and media

All experiments were performed at 20°C except where specified. All strains were maintained at 20°C, except strain FS428 *daf-2(e1370)* III (corresponding to the original CB1370 strain outcrossed 6 times) which was maintained at 19°C to prevent larval arrest. Strains were grown on nematode growth medium (NGM) agar plates freshly poured and seeded with *Escherichia coli* OP50 culture. The wildtype reference strain was *C. elegans* N2 Bristol. All strains used in this study are described in Table S1.

### Plasmids and generation of single-copy insertion alleles

The plasmids constructed for this study are described in Table S5. Plasmids used to create single copy insertion alleles by the miniMos method(Frøkjær-Jensen et al. 2014) are described in previous studies (Zhou et al. 2020, 2021) and the newly generated alleles are listed in Table S5. All constructs were verified by Sanger sequencing from GATC Company. For tissue specific expression, the promoters used were: myo-3– (body-wall muscle), unc-47– (GABAergic motoneuron), unc-17– (cholinergic motoneuron), rab-3– (pan-neuronal), dpy-7– (epidermis), or eft-3– (ubiquitous), N2 animals were injected with 15 ng/μl plasmid of interest containing the promoter and the open reading frame TIR1 or TOMM20 fused to fluorescent proteins, 50 ng/μl pCFJ601 (Mos1 transposase), 10 ng/μl pMA122 (negative selective marker Phsp16.2::peel-1), and 2.5 ng/μl pCFJ90 (Pmyo-2::mCherry). Neomycin (G418) was added to plates 24 h after injection at 1.5 μg/μl final concentration. Candidate plates were heat shocked for 2 h at 34°C. Selected lines were then bred to homozygosity and then crossed to generate the desired strains.

### Alleles generation by CRISPR/cas9 genome engineering

To generate the *kr462* allele, a flexible linker, the AID sequence(Zhang et al. 2015), another flexible linker and mNeonGreen were inserted tandemly into the *daf-2* locus, just before the stop codon. This sequence was PCR amplified with or without homology arms from pCR12 and the PCR products were used to generate the repair template (Dokshin et al. 2018). To generate the *kr535* allele, a flexible linker, the wrmScarlet, another flexible linker and 3 MYC tags were inserted tandemly into the *daf-16* locus, just before the stop codon. This sequence was PCR amplified with or without homology arms and the PCR products were used to generate the repair template(Dokshin et al. 2018). CrRNA were designed on *Benchling.com* and synthesized by IDT (Integrated DNA Technologies). For *kr462* TGAAAATGAGCATCTAATCG and ttttgggggttTCAGACAAG crRNA were used in tandem, for *kr535* CATGAGCTGAGTCAAGCTGG and tctctttcgaacaacaccag were used in tandem. The injection mix contained annealed double-stranded DNA donor cocktail as repair template at 200 ng/μl, Cas9 nuclease at 0.25 μg/μl (Integrated DNA Technologies), tracrRNA-crRNA duplex at 9 μM, pRF4 *[Peft-3::rol-6]* coinjection marker at 2.5 ng/μl, and RNase/DNase-free water qsp 20 μl. The candidate F1 animals were isolated by tracking the initial fluorescence knock-in in plates with F1 roller progeny. The F2 progenies were then isolated and homozygosed. The insertion was then confirmed by PCR and sequencing. Candidates were then outcrossed once with N2 before further crosses to generate the desired strains (See Table S1).

### RNAi

Bacterial feeding RNAi experiments were carried out essentially as described previously (Masse et al. 2008). Briefly, single colonies of bacteria containing plasmids of interest were first grown overnight in LB with 100 mg/ml ampicillin and 12.5 mg/ml tetracyclin and then for about 5 h in LB with 100 mg/ml ampicillin until OD600 reached 0,6 to 0,8. RNAi bacterial cultures were concentrated 6 times in order to increase their efficacy. Bacteria were seeded onto NGM plates containing 2 mM IPTG, 25 mg/ml carbenicillin and 1mM auxin. Worms were maintained on OP50 plates and transferred on RNAi plates at the L4 stage. The control (HT115) and D1081.2 *(unc-120)* clones were purchased from GeneService. Each clone has been sequenced to confirm its identity. In all RNAi experiments *rrf-3(pk1426)* mutants worms were used in order to increase RNAi sensitivity (Simmer *et al*. 2003).

### Ageing cohorts and auxin treatment

Auxin plates were prepared by adding auxin indole-3-acetic acid (Sigma-Aldrich) from a 400 mM stock solution in ethanol into NGM at the final concentration of 1 mM (Zhang et al. 2015). For control ethanol plates, the same volume of ethanol was added to NGM. Animals were transferred on auxin or ethanol plates at the L4 stage except for dauer tests for which eggs were laid and grown on auxin or ethanol plates.

For all ageing cohorts, 20 μM 5-fluorouracil (5-FU, Sigma-Aldrich) was added to prevent progeny growth. Animals were transferred weekly to fresh plates, without 5-FU after two weeks. The day of transition to L4 is counted as day 0 of adulthood for the cohort.

### Dauer and Lifespan assays

For dauer assays, L4s were transferred to ethanol or 1 mM auxin plates and allowed to mature into egg-laying adults, then removed after twenty-four hours. Progeny were assessed after forty-eight hours at 25°C for the presence of dauer larvae.

For lifespan assays, cohorts were handled as described in “ageing cohorts”. Lifespans were conducted at 20°C and were assessed every 2–3 days as described previously (Masse et al. 2008). Briefly, animals were scored as dead when they stopped moving and responding to prodding, they were censored when they crawled off the plate, had a “protruding vulva” or an “exploded vulva”. For oxidative stress survival assays, animals were transferred to 1 mM auxin plates at the L4 stage. After seven hours, young adult animals were transferred to 1 mM auxin plates supplemented with 20 mM paraquat (Methyl viologen dichloride hydrate, Sigma-Aldrich). Animals were scored as in lifespan assays but twice a day. All lifespan tests were blinded to avoid bias in the evaluation.

### Thrashing assays

For body bends frequency (BBF) measurements, 1-day-old or 13-day-old worms prepared as described in “ageing cohorts” were gently transferred into a 12-wells cell culture plate (ten worms per well) containing 1.5 mL of 2 % agarose in M9 buffer (3 g KH2PO4, 6 g Na2HPO4, 5 g NaCl, 0.25 g MgSO4·7 H2O, and distilled water up to 1 liter) and 2 mL of M9 buffer per well. Two minutes after transfer, animal movements were recorded for 30 s. BBF was then quantified using the open-source wrMTrck plugin on Fiji (ImageJ (v2.0.0)) software.

### Aldicarb assay

The tests were performed with synchronised worms. Twenty worms were added to a 100μl drop of 250μM aldicarb (Sigma, ref. 33386) in M9 buffer on polylysine-coated slides and incubated in a humidified chamber. 4, 4.5, 5 and 5.5 hours later, the worms were stimulated with a blue light for 50 seconds and touched with a worm pick. With the blue light still on, their movement was then recorded for 30 seconds with the same device as for the thrashing tests. The recordings were made blind and the worms were considered paralyzed if they made less than three body bends during this recording. Three biological replicates with independent cohorts were performed for each age.

### Microscopy and image processing

For all observations, animals were mounted on 2 % agarose dry pads with 4 % polystyrene beads (Polybeads, Polysciences) in M9 buffer. For images corresponding to **Figure 1a, 2, 5 and S1**, worms were observed using an Andor spinning disk system (Oxford Instruments) installed on a Nikon-IX86 microscope (Olympus) equipped with a 40x/NA 1.3 and a 60×/NA 1.42 oil-immersion objectives and an Evolve electron-multiplying charge-coupled device camera. Each animal was imaged with IQ software (APIS Informationstechnologien) as a stack of optical sections (0.3 μm apart) across the whole thickness of the worm (Figure 1a and S1a) or across a specific tissue **(Figure 2 and 4).** All images were processed using the Fiji (ImageJ) software and correspond to the sums of the same number of slices (for each strain with or without auxin) except for images of the intestine (**Figure 2g and S1h)** which correspond to z projection of maximum intensity. For fluorescent quantification images covering the whole head including the most posterior pharyngeal bulb were summed (85 slices) and a ROI (180 × 230 pixels) was positioned relative to the middle of the most posterior part of the pharynx.

### Scoring of DAF-16 nuclear localization

One-day-old adults were observed using a AZ100 Multizoom Microscope (Nikon) equipped with a CMOS flash 4 C11440 (Hamammatsu) camera, for no more than 10 minutes per slide to avoid post-mounting DAF-16::wSCARLET nuclear translocation. In absence of auxin, the fluorescence associated with DAF-16::wSCARLET appeared diffuse in the cytoplasm and nuclei of most tissues, except in the head neurons where it was completely excluded from the nuclei. A tissue was scored positive when several nuclei were brighter than the cytoplasm. Strains scoring and image analysis were performed blind.

### Scoring of muscle mitochondria morphology

Images of *krSi134*[Pmyo-3::tom-20N::wScarlet] worms were acquired on an Axioscop compound microscope (Zeiss) equipped with a Neofluar 63x/NA 1.25 oil-immersion objective and a EMCC CoolSnap HQ (Photomectrics) camera. For each worm a representative image from the posterior body wall muscle cells was acquired. Cells with long interconnected mitochondrial networks were classified as interconnected; cells with a combination of interconnected mitochondrial networks along with some smaller fragmented mitochondria were classified as interrupted; cells with sparse small round mitochondria were classified as fragmented. Strains scoring and image analysis were performed blind.

### Statistical analysis

All statistical analyses were performed with R version 4.0.1 (2020-06-06). The R Test Survival Curve Differences (package survival_3.2-3) was used to analyze lifespan assays. This test is based on the *G-rho* family of tests which makes use of the Kaplan-Meier estimate of survival(Fleming et al. 1982). Thrashing assays were analyzed using the non-parametric Kruskal-Wallis Rank Sum Test, followed by Dunn’s Test of Multiple Comparisons (package rstatix_0.6.0) with FDR adjusting method as post-hoc tests. To compare between different conditions for the mitochondria morphology assay a Fisher exact test was performed, followed by pairwise tests with FDR adjusting method as post-hoc tests (package rcompanion_2.4.1). For all tests compared samples were considered different when statistical test gave an adjusted P-value <0.05 (*p<0.05; **p<0.01; ***p<0.001), ns: non-significant.

## Acknowledgement

We thank the Caenorhabditis Genetic Center (which is funded by NIH Office of Research Infrastructure Programs, P40 OD010440) for strains. We are grateful to Laure Granger for technical assistance. This work was supported by the European Research Council (ERC_Adg C.NAPSE #695295), within the framework of the LABEX CORTEX (ANR-11-LABX-0042) of Université de Lyon operated by the French National Research Agency (ANR). CR was supported by a PhD fellowship from the French Ministry of Higher Education and Fondation de la recherche médicale (FDT202106012780). Biorender.com images were used to generate the original graphical abstract.

## Conflict of interest

The authors declare no conflict of interest.

## Authors contributions

F.S and C.R. conceived the study. C. R., L. M., A. A., M. S., and F. S. performed the experiments. B. B. and C. V. constructed the mitochondria reporter strain and the tir1 alleles respectively. F.S. and C.R. analyzed and interpreted the data. The manuscript was written by F.S. and C. R. and edited by LM and JLB.

## Data availability

The authors confirm that the data supporting the findings of this study are available within the article and its supplementary materials.

## Supporting information listing (Figures and Tables)

**Figure S1**: DAF-2::AID::mNG was efficiently degraded in different tissues in presence of auxin.

**Figure S2**: Survival curve of biological replicates corresponding to Table S2.

**Figure S3**: Ubiquitous DAF-2 depletion induced aldicarb hypersensitivity.

**Figure S4**: Muscle or neuronal depletion of DAF-2 was sufficient to prevent mitochondria fragmentation during ageing

**Table S1**: Strains description

**Table S2**: Lifespan data and statistics corresponding to lifespan curves shown in Figures 1, 3 and Figure S2.

**Table S3**: Regulation of dauer arrest in worms with tissue-specific depletion of DAF-2

**Table S4**: Numbers of worms with DAF-16::wrmSCARLET nuclear accumulation in different tissues

**Table S5**: Plasmids and alleles generated

---

**Supplementary Figure S1:**
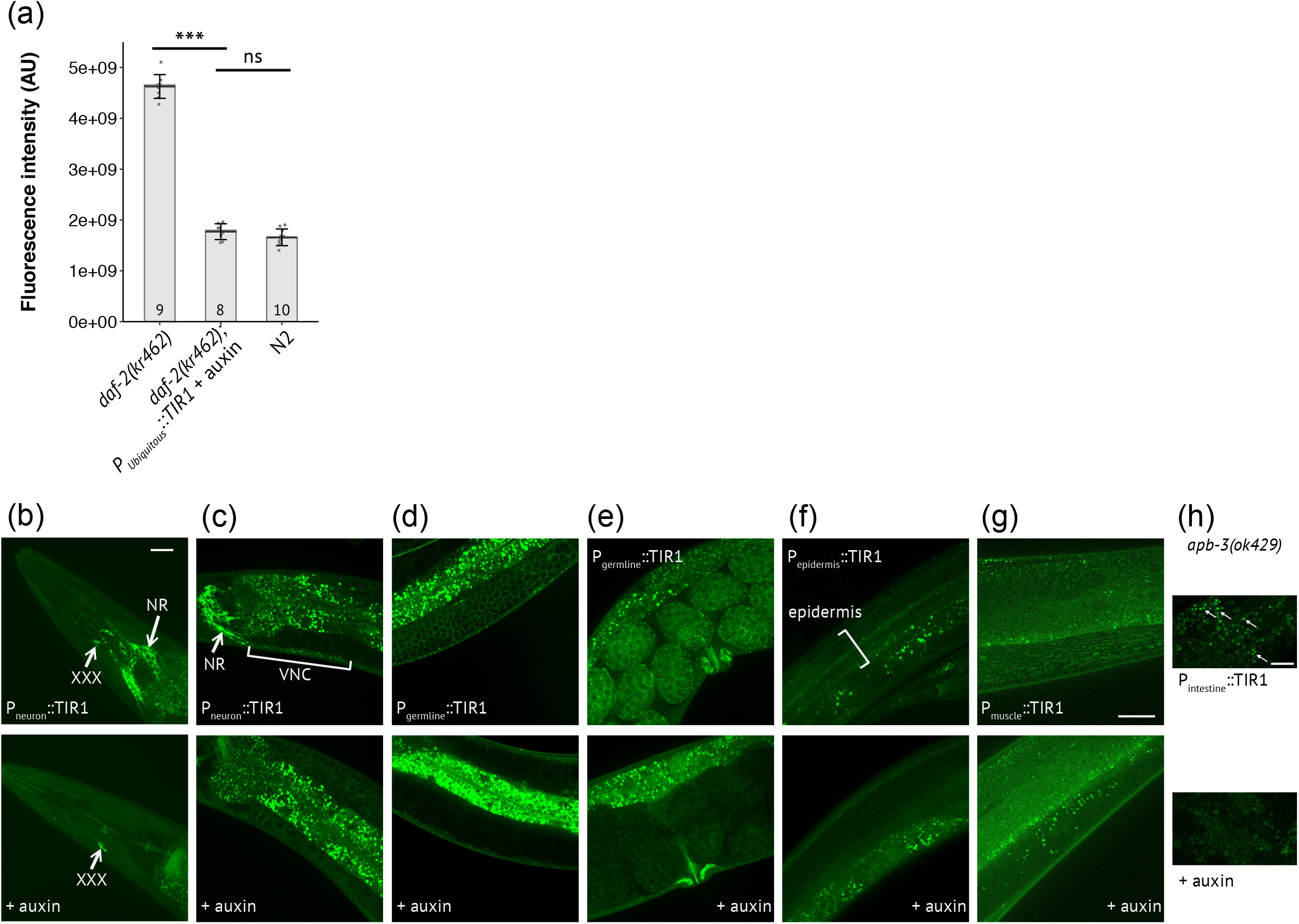
DAF-2::AID::mNG is efficiently degraded in different tissues in presence of auxin. (**a)** Quantification of fluorescence associated with DAF-2::AID::mNG expression in the absence or presence of 1mM auxin as indicated. Worms were transferred to auxin-containing plates at the L4 stage. Twenty-four hours later, the heads of the animals were imaged. and quantified (see Experimental procedures). N2 were used as control for nonspecific autofluorescence. The numbers of animals scored is indicated in bars. Bars indicate median values, means are represented by black horizontal lines and brackets show standard deviations, ns: non-significant, ***: p < 0.001, Kruskal-Wallis and Dunn’s post hoc test with FDR method for adjusting p-value. **(b-h)** Images of DAF-2::AID::mNG expression in 1-day-old animals. Animals express transgenic TIR1 under the control of tissue specific promoters, as indicated, in the absence (upper panel) or in the presence (lower panel) of 1 mM auxin for 24 hours. **(a-b)** With a neuronal promoter, DAF-2::AID::mNG was no longer detected in the nerve ring (NR) **(b)** and in the ventral nerve cord (VNC) **(c)** but was still detected in the XXX cell **(b)**. **(d,e)** With a germline promoter, DAF-2::AID::mNG signal was reduced in the proliferating germ cells **(d)** and in the eggs **(e)**. **(f)** With a epidermal promoter, DAF-2::AID::mNG disappeared from the epidermis syncytium. **(g)** With a muscular promoter, DAF-2::AID::mNG was no longer detected in body wall muscles. **(h)** With an intestinal promoter, DAF-2::AID::mNG signal was downregulated in the intestine. *apb-3(ok429)* mutants with reduced intestinal autofluorescence were used. Scale bars: 20 μm **(b-g)**, 10 μm **(h)**.

**Supplementary Figure 2:**
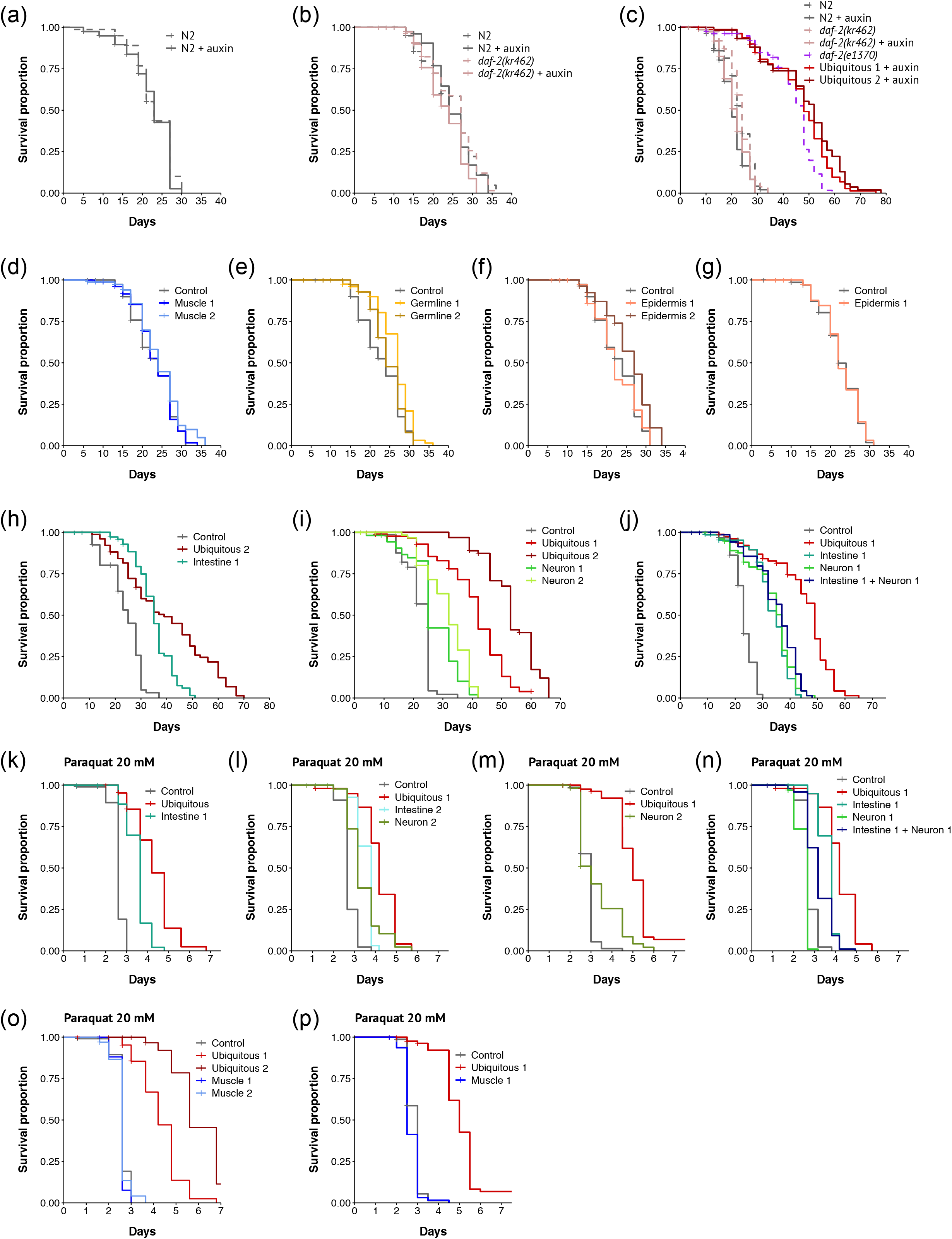
Survival curve of biological replicates corresponding to Table S2. Survival curves of animals with DAF-2 depletion in none, all cells or in the indicated tissue in absence **(a-j)** or in presence of 20 mM paraquat (**k-p).** All conditions corresponding to (**d-p**) were in presence of 1mM auxin. For detailed genotypes and statistics, see Table S2.

**Supplementary Figure 3:**
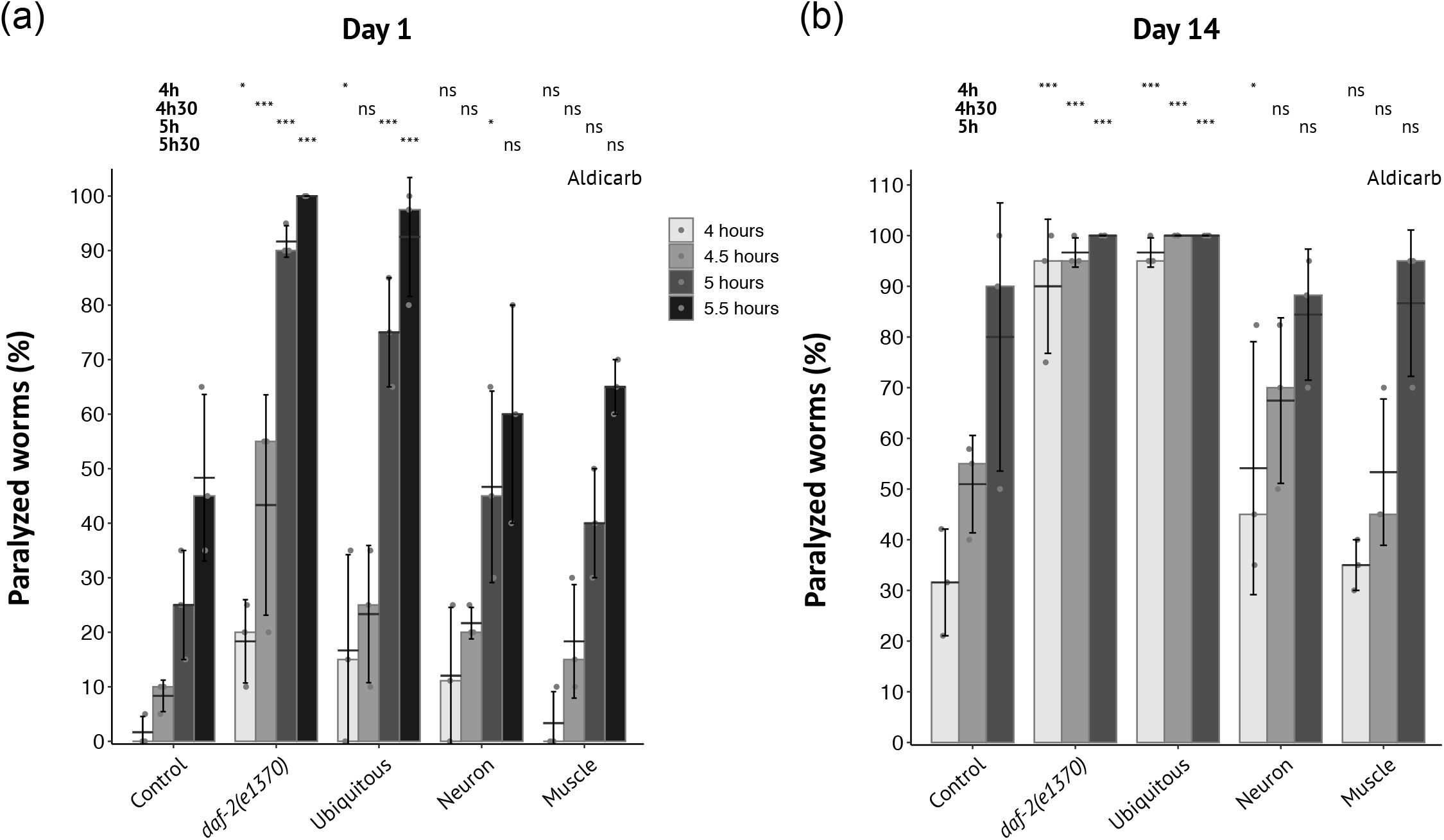
Ubiquitous DAF-2 depletion induced aldicarb hypersensitivity. Synchronized 1-day-old **(a)** or 14-day-old **(b)** adults with ubiquitous (ubiquitous 1), neuronal (neuron 2) or muscle (muscle 1) DAF-2 depletion from the L4 stage, or *daf-2(e1370)* mutants, as indicated, were placed in a drop of M9 containing 250 μM aldicarb. Control corresponds to *daf-2(kr462)* worms cultivated in presence of auxin from the L4 stage. The cholinesterase inhibitor aldicarb causes an accumulation of acetylcholine in the synaptic cleft of neuromuscular junctions, resulting in sustained muscle activation and eventually paralysis. Worms were assayed for paralysis over time (see Experimental procedure). Data from 3 independent experiments have been pooled (n= 60 individuals per condition). Comparisons used Fisher exact test, followed by pairwise tests with FDR adjusting method as post-hoc tests: ns: not significant, *: p_adjusted_ < 0.05, ***: p_adjusted_ < 0.001. Statistics are presented in several lines corresponding to the comparison with the control strain at each time point.

**Supplementary Figure 4:**
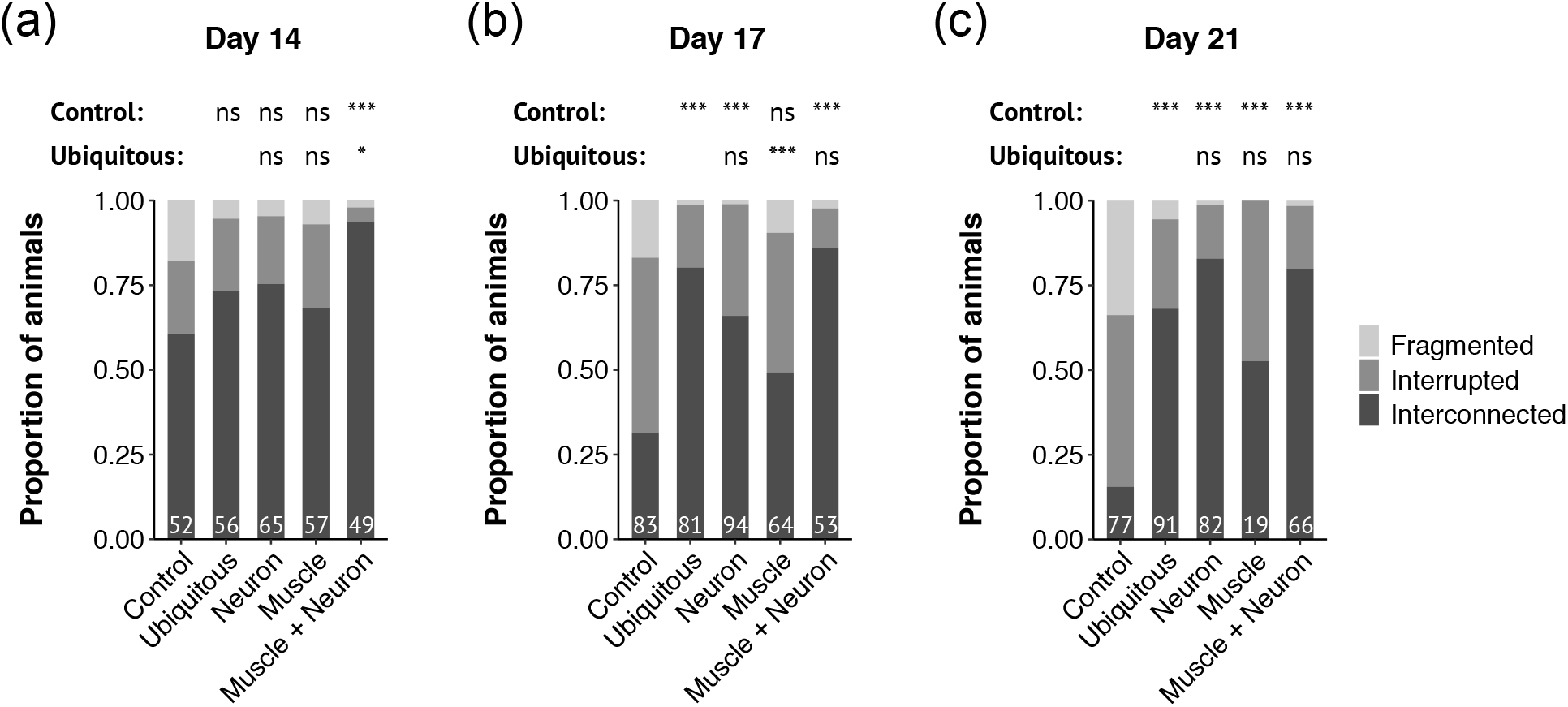
Muscle or neuronal depletion of DAF-2 is sufficient to prevent muscle mitochondria fragmentation during ageing. **(a-c)** Quantification of muscular mitochondrial morphology at day 14 **(a)**, day 17 **(b)**, or day 21 **(c)** of adulthood in worms with DAF-2 depletion in all tissues, or in muscle, neurons or muscle and neurons as indicated. The number of animals scored is indicated in each bar. Data correspond to a pool of 3 to 5 independent trials depending on the strain. Strains used were: *daf-2(kr462)* in presence of auxin as Control, Ubiquitous 1, Neuron 1 and 2, Muscle 1, Muscle 1 + Neuron 1 (see Table S1 for strain description). Comparisons used Fisher exact test, followed by pairwise tests with FDR adjusting method as post-hoc tests: ns: not significant, *: p_adjusted_ < 0.05, ***: p_adjusted_ < 0.001. Statistics are presented as two lines that include comparison with the control and Ubiquitous 1 strain.

**Table S1:**
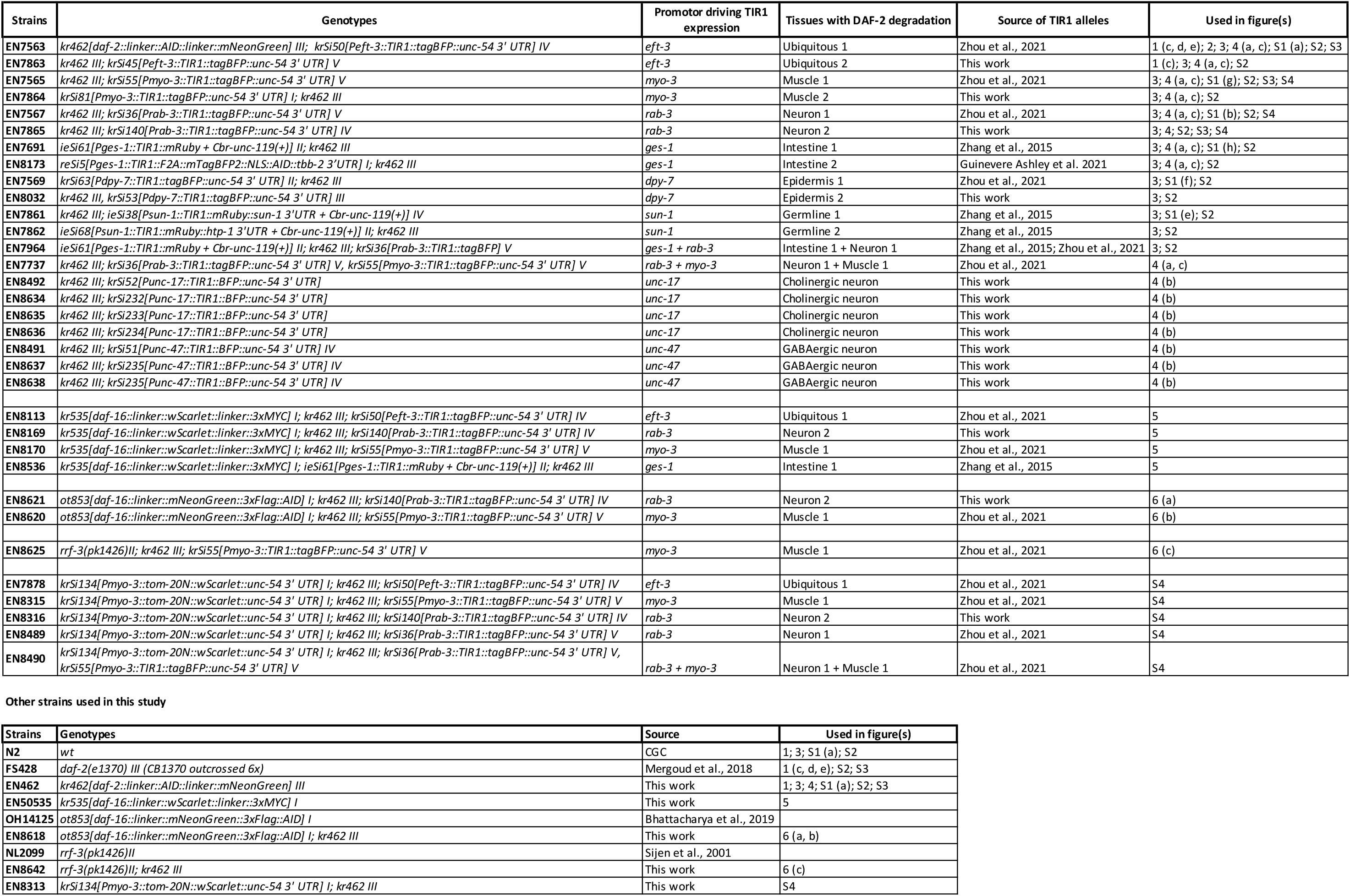
Strains.

**Table S2:**
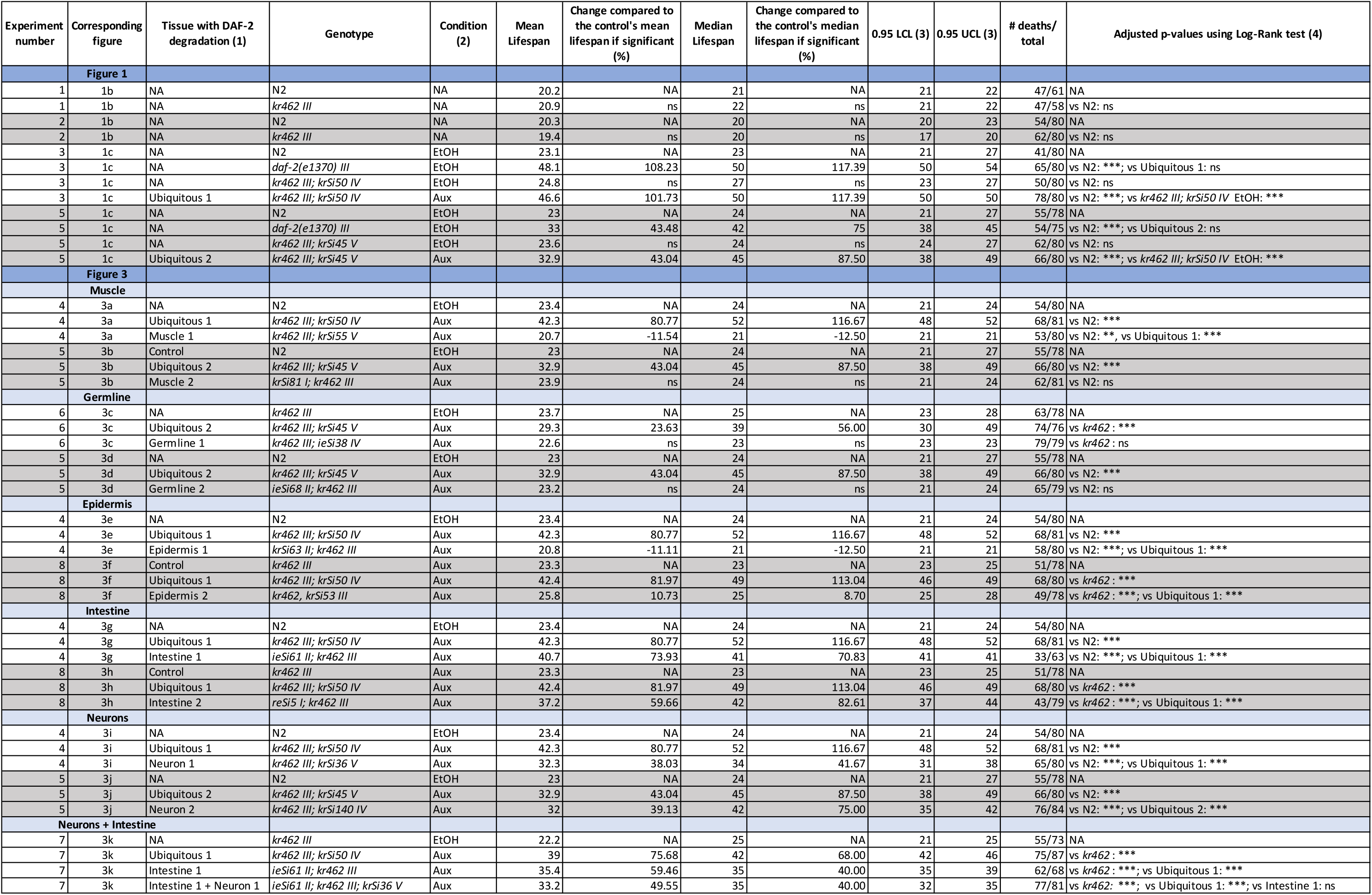
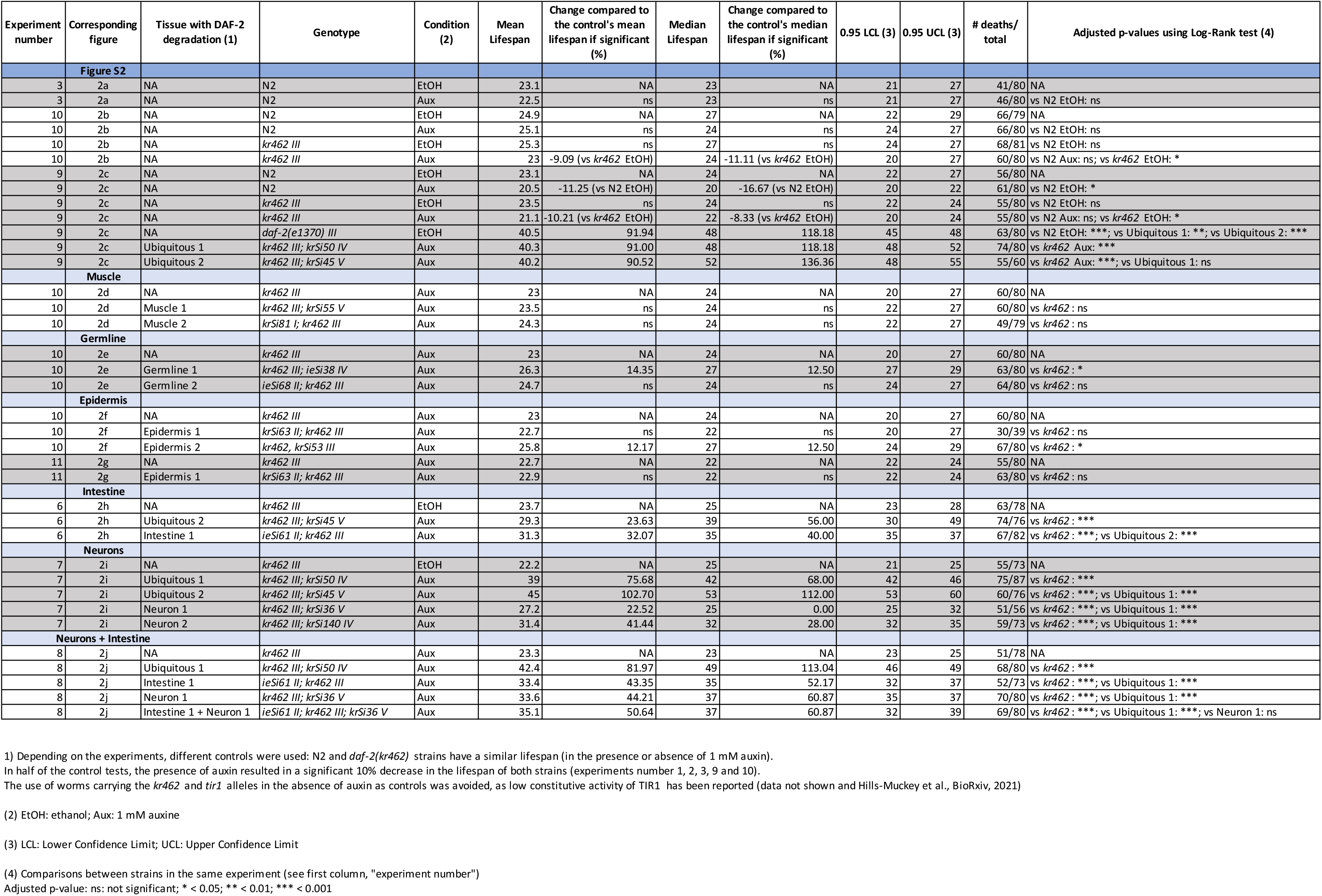
Lifespan data and statistics corresponding to lifespan curves shown in Figures 1, 3 and S2.

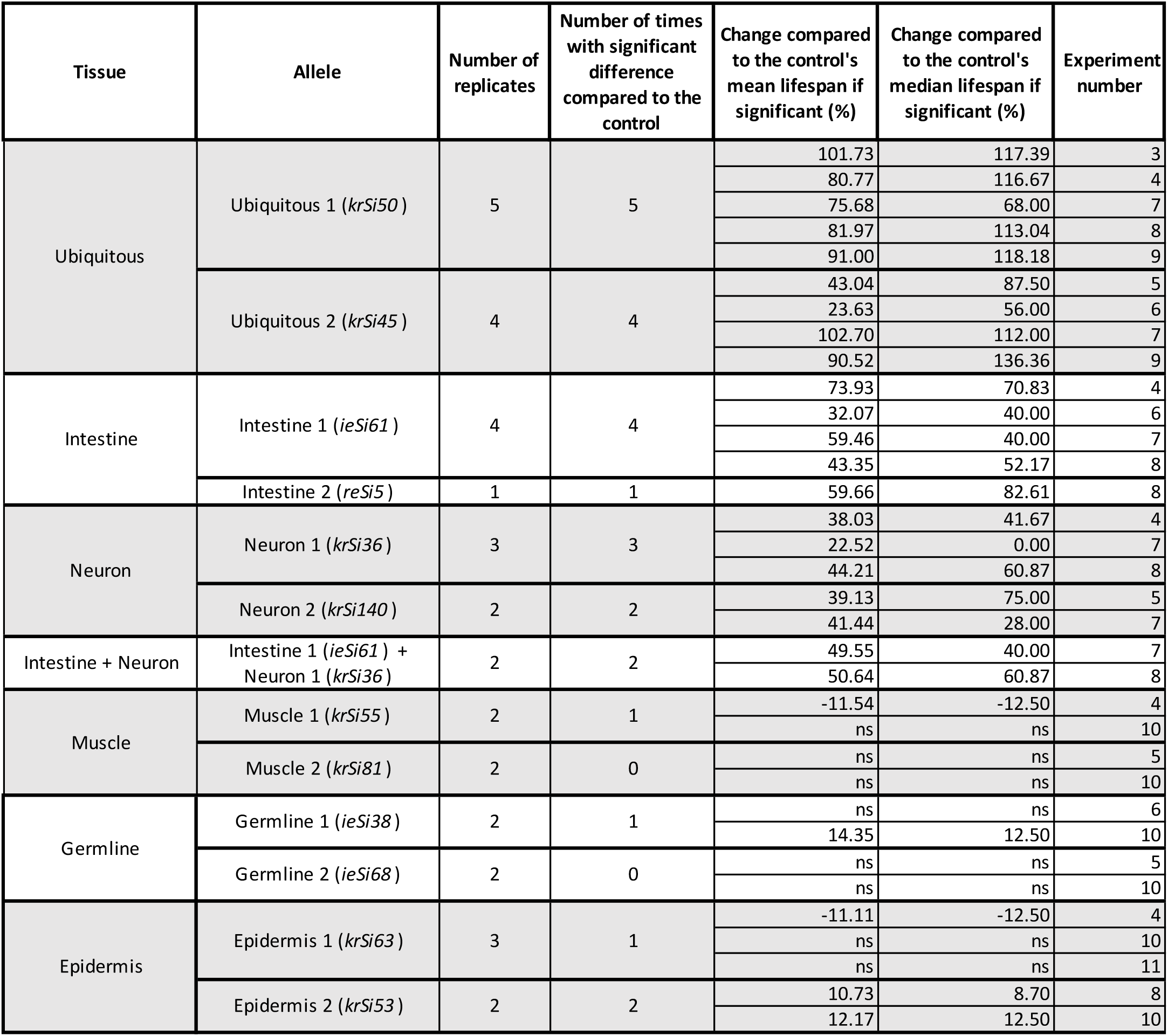
Summary of lifespan replicates per strain and results.

**Table S2:**
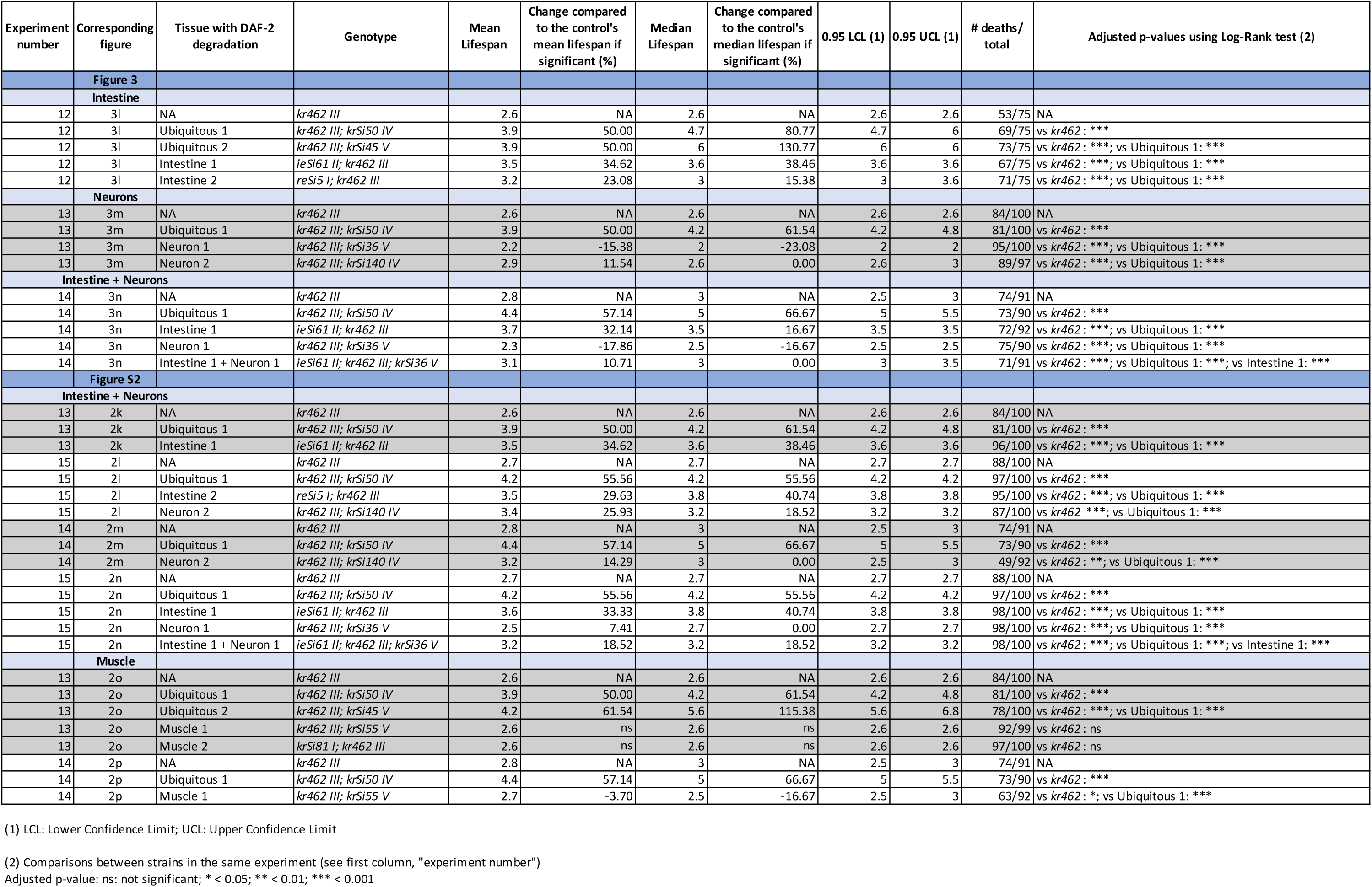
Lifespan data and statistics corresponding to lifespan curves in presence of paraquat and auxine shown in Figures 3 and S2.

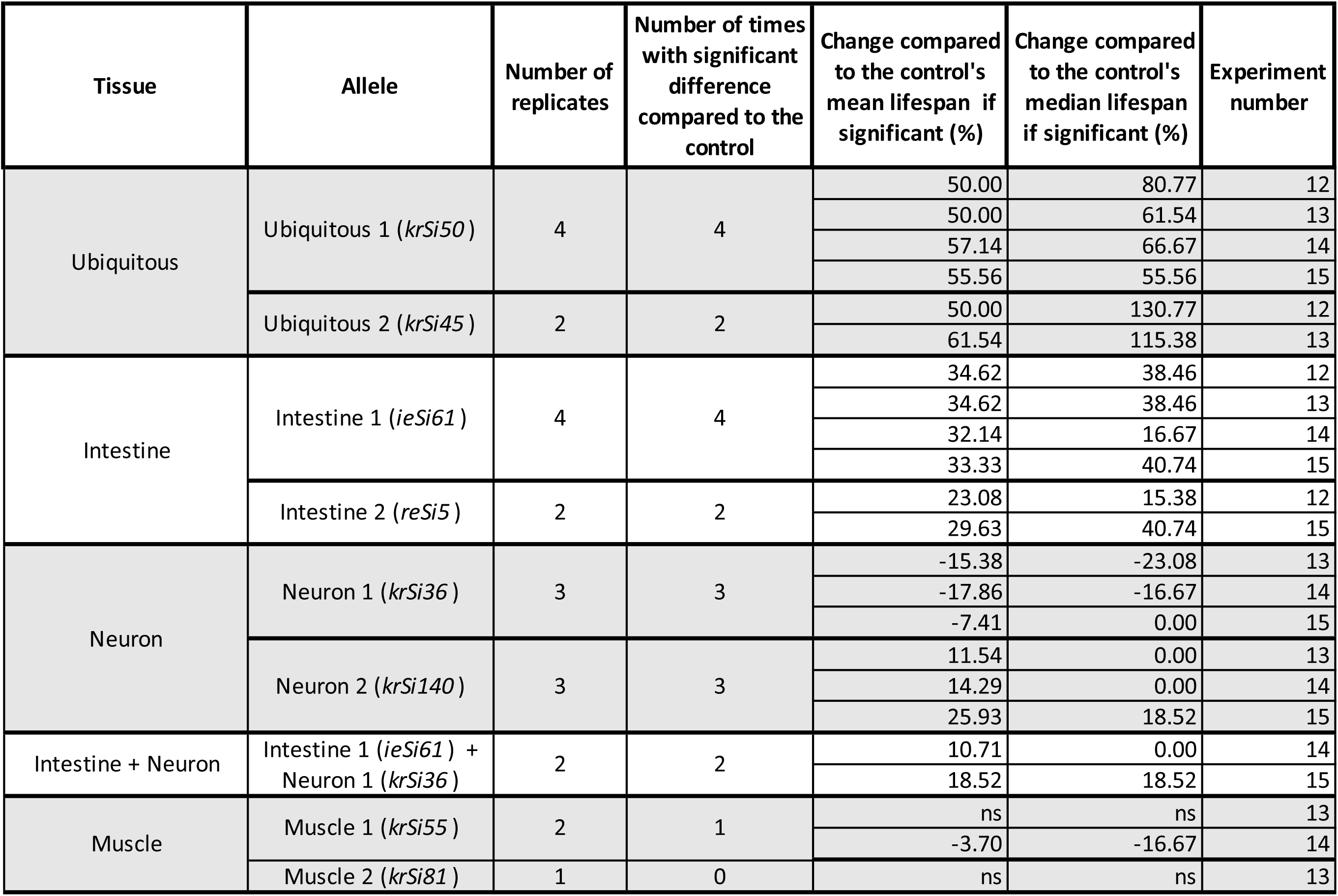
Summary of lifespan replicates in presence of paraquat per strain and results.

**Table S3:**
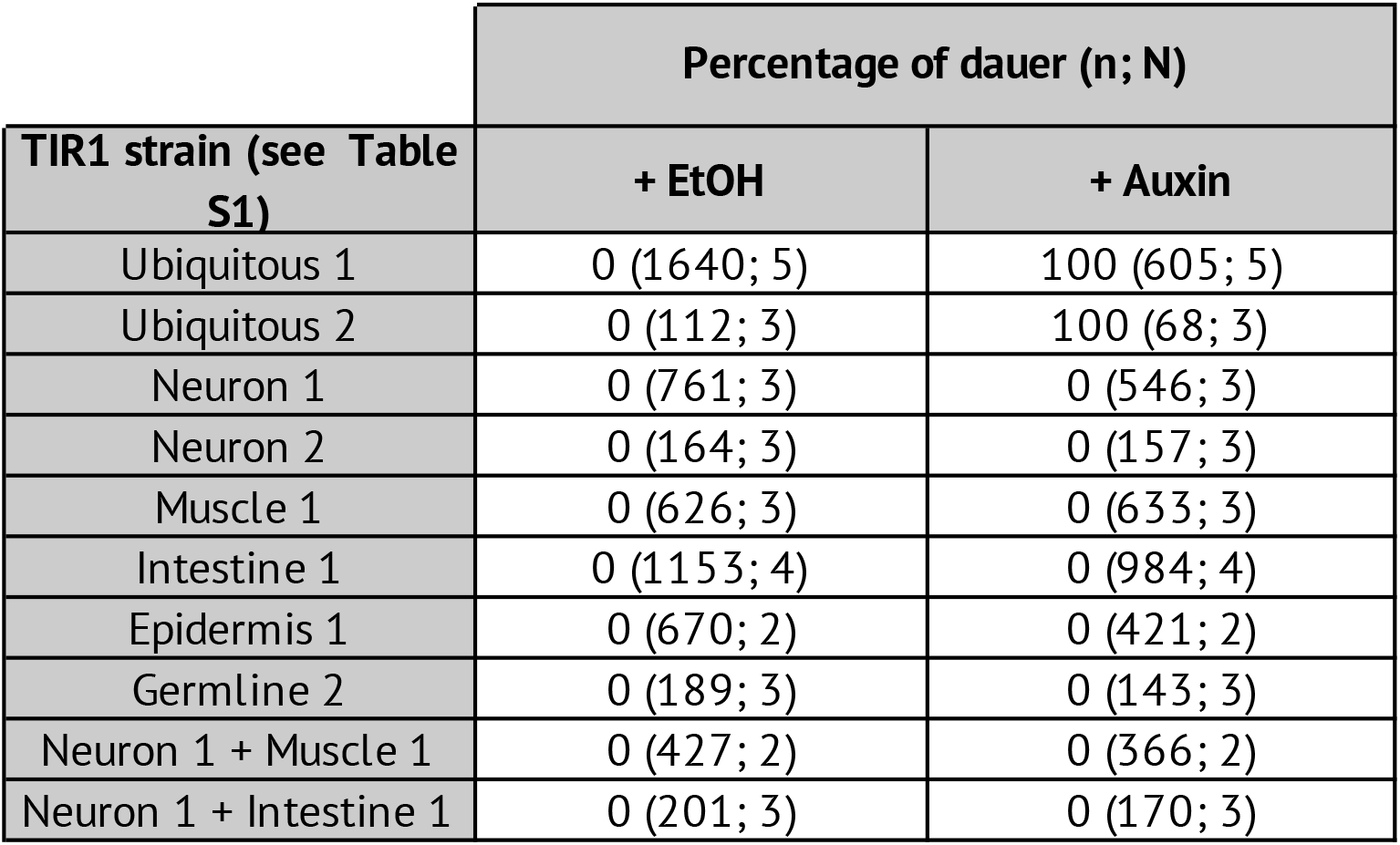
dauer percentage.

| **TIR1 strain (see Table S1)** | **Percentage of dauer (n; N)** |  |
| --- | --- | --- |
|  | **+ EtOH** | **+ Auxin** |
| Ubiquitous 1 | 0 (1640; 5) | 100 (605; 5) |
| Ubiquitous 2 | 0 (112; 3) | 100 (68; 3) |
| Neuron 1 | 0 (761; 3) | 0 (546; 3) |
| Neuron 2 | 0 (164; 3) | 0 (157; 3) |
| Muscle 1 | 0 (626; 3) | 0 (633; 3) |
| Intestine 1 | 0 (1153; 4) | 0 (984; 4) |
| Epidermis 1 | 0 (670; 2) | 0 (421; 2) |
| Germline 2 | 0 (189; 3) | 0 (143; 3) |
| Neuron 1 + Muscle 1 | 0 (427; 2) | 0 (366; 2) |
| Neuron 1 + Intestine 1 | 0 (201; 3) | 0 (170; 3) |

**Table S4:** DAF-16 nuclear translocation.

|  |  | Percentage of worms with strong DAF-16::wSCARLET nuclear signal (n) |  |  |
| --- | --- | --- | --- | --- |
|  |  | Intestine | Neurons | Muscles |
| DAF-2 degradation | Control | 0 (22) | 0 (25) | 0 (22) |
|  | Ubiquitous | 100 (31) | 100 (28) | 100 (32) |
|  | Intestine | 100 (21) | 0 (21) | 100 (22) |
|  | Neuron | 0 (25) | 100 (28) | 0 (25) |
|  | Muscle | 0 (25) | 0 (25) | 100 (25) |

**Table S5.**
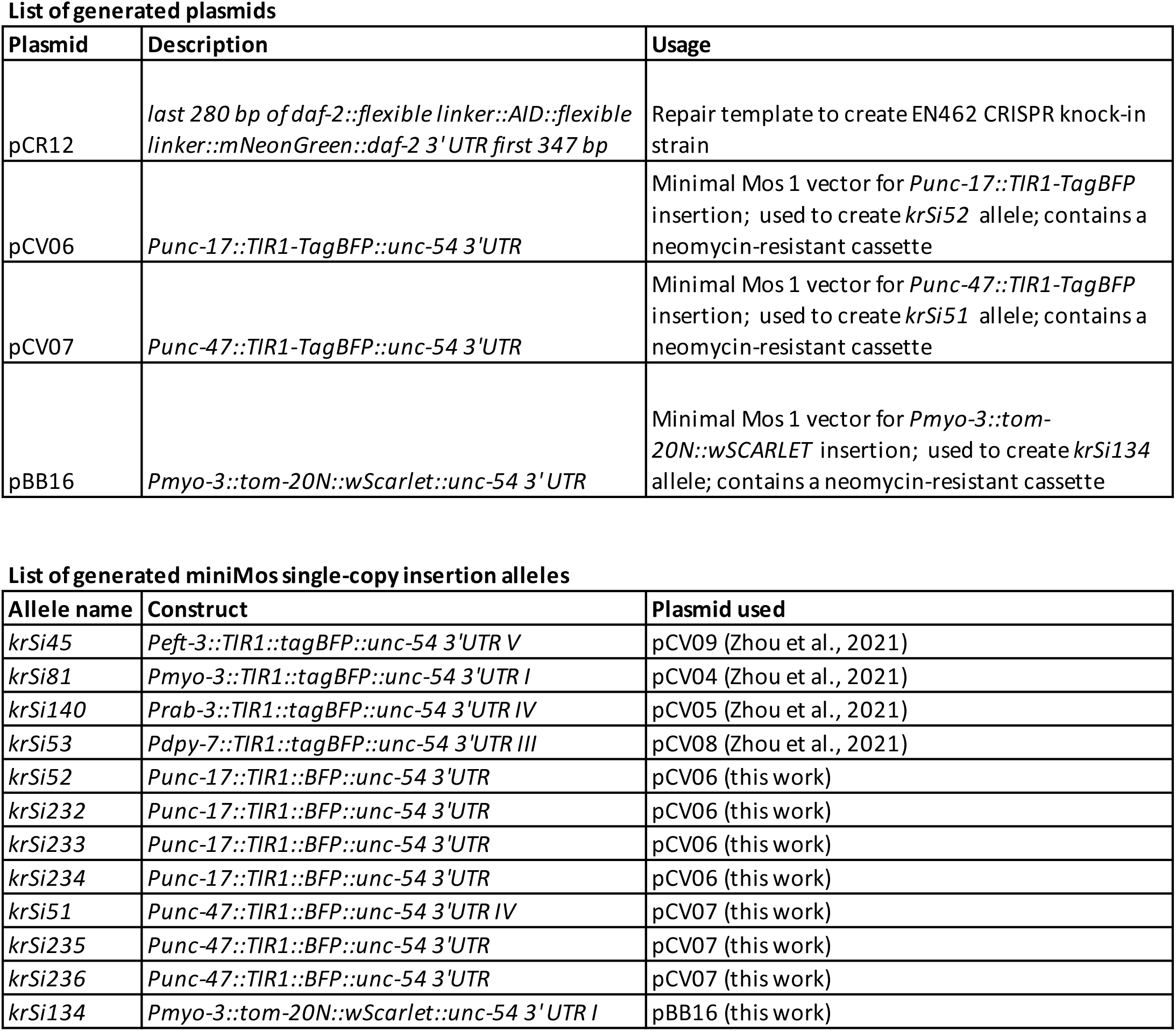
List of generated plasmids

| Plasmid | Description | Usage |
| --- | --- | --- |
| pCR12 | *last 280 bp of daf-2::flexible linker::AID::flexible linker::mNeonGreen::daf-2 3' UTR first 347 bp* | Repair template to create EN462 CRISPR knock-in strain |
| pCV06 | *Punc-17::TIR1-TagBFP::unc-54 3' UTR* | Minimal Mos 1 vector for *Punc-17::TIR1-TagBFP* insertion; used to create *krSi52* allele; contains a neomycin-resistant cassette |
| pCV07 | *Punc-47::TIR1-TagBFP::unc-54 3' UTR* | Minimal Mos 1 vector for *Punc-47::TIR1-TagBFP* insertion; used to create *krSi51* allele; contains a neomycin-resistant cassette |
| pBB16 | *Pmyo-3::tom-20N::wScarlet::unc-54 3' UTR* | Minimal Mos 1 vector for *Pmyo-3::tom-20N::wSCARLET* insertion; used to create *krSi134* allele; contains a neomycin-resistant cassette |

List of generated miniMos single-copy insertion alleles
| Allele name | Construct | Plasmid used |
| --- | --- | --- |
| *krSi45* | *Peft-3::TIR1::tagBFP::unc-54 3' UTR V* | pCV09 (Zhou et al., 2021) |
| *krSi81* | *Pmyo-3::TIR1::tagBFP::unc-54 3' UTR I* | pCV04 (Zhou et al., 2021) |
| *krSi140* | *Prab-3::TIR1::tagBFP::unc-54 3' UTR IV* | pCV05 (Zhou et al., 2021) |
| *krSi53* | *Pdpy-7::TIR1::tagBFP::unc-54 3' UTR III* | pCV08 (Zhou et al., 2021) |
| *krSi52* | *Punc-17::TIR1::BFP::unc-54 3' UTR* | pCV06 (this work) |
| *krSi232* | *Punc-17::TIR1::BFP::unc-54 3' UTR* | pCV06 (this work) |
| *krSi233* | *Punc-17::TIR1::BFP::unc-54 3' UTR* | pCV06 (this work) |
| *krSi234* | *Punc-17::TIR1::BFP::unc-54 3' UTR* | pCV06 (this work) |
| *krSi51* | *Punc-47::TIR1::BFP::unc-54 3' UTR IV* | pCV07 (this work) |
| *krSi235* | *Punc-47::TIR1::BFP::unc-54 3' UTR* | pCV07 (this work) |
| *krSi236* | *Punc-47::TIR1::BFP::unc-54 3' UTR* | pCV07 (this work) |
| *krSi134* | *Pmyo-3::tom-20N::wScarlet::unc-54 3' UTR I* | pBB16 (this work) |

## Notes

### Competing Interest Statement

The authors have declared no competing interest.

